# An apical junction protein antagonizes mechanosensitive calcium signaling to establish stochastic choices of olfactory neuron subtypes

**DOI:** 10.64898/2026.03.12.710951

**Authors:** Rui Xiong, Jun Yang, Shengyao Yuan, Eleana Liu, Xiaohong Wang, Chiou-Fen Chuang

**Author notes:** Corresponding author: Chiou-Fen Chuang.

## Abstract

Mechanical forces regulate brain development and left-right body patterning. However, the role of mechanical signaling in brain lateralization remains unclear. In *Caenorhabditis elegans*, the left and right AWC olfactory neurons communicate via a gap junction network to stochastically differentiate into the default AWC^OFF^ and induced AWC^ON^ subtypes. SLO BK potassium channels, SLO-1 and SLO-2, act redundantly to inhibit a calcium-regulated protein kinase pathway in the specification of the AWC^ON^ subtype. Here, we identified a role for AJM-1 (apical junction molecule 1) in promoting AWC^ON^ from an unbiased forward genetic screen for mutants that suppress the *slo-1(gf)* 2AWC^ON^ phenotype. AJM-1 is located at three distinct tight junctions between amphid neurons (including AWC) and sheath glia, sheath and socket glia, and socket glia and hypodermal cells (also known as epidermal cells) at the anterior tip of the animal. In addition to its cell-autonomous function, the non-cell-autonomous function of AJM-1 in glial and hypodermal cells is required for the specification of the AWC^ON^ subtype. Furthermore, we identified a role for the DEL-1 mechanosensitive DEG/ENaC channel in the calcium signaling pathway, mediated by UNC-2 and EGL-19 voltage-activated calcium channels, that specifies AWC^OFF^. Together, our results suggest a mechanism in which AJM-1 promotes SLO-1 expression and antagonizes mechanosensitive calcium signaling, thereby promoting the AWC^ON^ subtype. This study provides insight into the role of mechanical force in the stochastic lateralization of olfactory neurons.

## Introduction

Mechanical forces have been implicated in brain development (Javier-Torrent, Zimmer-Bensch et al. 2021). Mechanotransduction is mediated by mechanosensitive proteins, including the degenerin/epithelial sodium channel (DEG/ENaC) superfamily, transient receptor potential channel (TRP) family, two-pore domain potassium channel (K2P) family, and Piezo ion channel family (Otero-Sobrino, Blanco-CarlÛn et al. 2023), which allow for physical forces to be sensed and transformed into biochemical signals. In the nervous system, migrating and differentiating neurons experience mechanical forces from interactions with nearby cells and the extracellular matrix, leading to mechanotransduction that regulates critical processes (Javier-Torrent, Zimmer-Bensch et al. 2021). For example, ligand-receptor interactions generate a mechanical force that activates DEG/ENaC channels to drive the influx of calcium through L-type voltage-activated calcium channels, thereby stimulating further growth of dendritic branches in *Caenorhabditis elegans* (Tao, Coakley et al. 2022).

Although mechanical forces have been shown to function in the left-right patterning of the body (Ferreira and Vermot 2017, Du, Li et al. 2023), the role of mechanical forces in brain lateralization is not determined. In *C. elegans*, the left and right amphid wing “C” (AWC) olfactory neuron pair asymmetrically differentiates into two subtypes, the default AWC^OFF^ and induced AWC^ON^, in a stochastic manner during embryogenesis (Troemel, Sagasti et al. 1999, Alqadah, Hsieh et al. 2016). AWC neurons are located in the primary sense organs, the amphid sensilla, each of which consists of 11 chemosensory neurons, one thermosensory neuron, one sheath glial cell, and one socket glial cell in the head (Ward, Thomson et al. 1975, Bae 2008, Singhvi and Shaham 2019). Previous studies demonstrated three distinct electron-dense tight junctions between amphid neurons and sheath glia, between sheath and socket glia, and between socket glia and hypodermal cells at the anterior tip of the animal (Ward, Thomson et al. 1975, Perkins, Hedgecock et al. 1986, Low, Williams et al. 2019, Singhvi and Shaham 2019). In the anterior sensilla (including the amphid sensilla), soma of developing neurons (including AWC) and glial cells migrate posteriorly from anchored dendrites and glial processes, respectively, toward their final destinations via retrograde extension (Heiman and Shaham 2009, Kelley, Yochem et al. 2015, Singhvi and Shaham 2019). The inward-pulling mechanical force exerted by AWC and glial somas during retrograde extension may modulate mechanical tension at the tight junctions between AWC and sheath glia (and perhaps between sheath and socket glia, and between socket glia and hypodermal cells), thereby regulating the choice of asymmetric AWC subtypes. Therefore, AWC neuronal asymmetry provides a unique model for studying the potential role of mechanical force in the stochastic lateralization of the nervous system.

The AWC neuron pair exhibits asymmetry at both the molecular and functional levels. The AWC^ON^ neuron expresses the chemoreceptor *str-2* and detects butanone, while the AWC^OFF^ neuron expresses *srsx-3* and senses 2,3-pentane-dione (Troemel, Sagasti et al. 1999, Wes and Bargmann 2001, Bauer Huang, Saheki et al. 2007). Additionally, AWC asymmetry is stochastic in that the left and right AWC neurons have equal probabilities of acquiring either subtype (Troemel, Sagasti et al. 1999, Hsieh, Alqadah et al. 2014, Alqadah, Hsieh et al. 2016). Calcium plays a dual role in AWC asymmetry: autonomously promoting the default AWC^OFF^ subtype and non-autonomously promoting the induced AWC^ON^ subtype (Schumacher, Hsieh et al. 2012). The AWC^OFF^ subtype is specified by a calcium-activated protein kinase pathway. In this pathway, calcium influx through voltage-activated calcium channels (comprised of the α1 subunits UNC-2 N-type or EGL-19 L-type and the α2δ subunit UNC-36) activates a downstream calcium-signaling complex consisting of UNC-43 calcium/calmodulin-dependent protein kinase (CaMKII), TIR-1 (SARM1) adaptor protein, and NSY-1 MAP kinase kinase kinase (ASK1 MAPKKK) (Troemel, Sagasti et al. 1999, Sagasti, Hisamoto et al. 2001, Chuang and Bargmann 2005, Bauer Huang, Saheki et al. 2007, Chang, Hsieh et al. 2011). Intercellular calcium signaling through a transient NSY-5 gap junction neural network, including AWC and at least 17 non-AWC neuron pairs, non-cell autonomously coordinates a precise 1AWC^ON^/1AWC^OFF^ choice (Chuang, Vanhoven et al. 2007, Schumacher, Hsieh et al. 2012). Voltage- and calcium-activated SLO BK potassium channels act downstream of the NSY-5 gap junction protein and NSY-4 claudin-like protein, to antagonize calcium signaling in the AWC^ON^ neuron (Alqadah, Hsieh et al. 2016). However, the mechanism underlying calcium signaling suppression in the AWC^ON^ subtype is not fully understood.

Here, we identified an <u>a</u>pical junction <u>m</u>olecule (AJM-1) that promotes the AWC^ON^ subtype from an unbiased forward genetic screen for mutants that suppress the *slo-1* gain-of-function 2AWC^ON^ phenotype. AJM-1 protein is broadly expressed in the junction of multiple tissues, including AWC neurons, glia, and hypodermal cells. We show that AJM-1 activity in non-neuronal cells, including glial and hypodermal cells, is sufficient and required for the AWC^ON^ neuron subtype specification. In addition, *slo-1* expression is reduced in AWC cells in *ajm-1* loss-of-function mutants. Our genetic data suggest that DEG/ENaC mechanosensitive channels promote the AWC^OFF^ subtype by activating voltage-activated calcium channels. Furthermore, AJM-1 may antagonize the activity of DEG/ENaCs, thereby suppressing calcium signaling and promoting the AWC^ON^ subtype. Together, our study suggests a model in which AJM-1 functions cell autonomously and non-cell autonomously to promote SLO-1 expression and antagonize the activity of DEG/ENaC mechanosensitive channels, probably by regulating the mechanical tension between non-neuronal and neuronal cells, thereby inhibiting calcium signaling in the AWC^ON^ neuron.

## Results

### The *vy11* mutation causes a defect in AWC asymmetry

In wild-type animals, the AWC^ON^ neuron expresses the G protein-coupled receptor (GPCR) gene *str-2* (Troemel, Sagasti et al. 1999), while the contralateral AWC^OFF^ neuron expresses the GPCR gene *srsx-3* (Bauer Huang, Saheki et al. 2007), resulting in the 1AWC^ON^/1AWC^OFF^ phenotype (Figures 1Ai, 1B, rows a and n, and S1A). *slo-1(ky399)* gain-of-function (gf) mutants lost expression of the AWC^OFF^ marker *srsx-3* and expressed the AWC^ON^ marker *str-2* in both AWC cells, resulting in the 2AWC^ON^ phenotype (Alqadah, Hsieh et al. 2016) (Figures 1Aii and S1A). We identified the *vy11* allele from an unbiased forward genetic screen for mutants that suppressed the *slo-1(ky399gf)* 2AWC^ON^ phenotype. *vy11* mutants lost expression of the AWC^ON^ marker *str-2* and expressed the AWC^OFF^ marker *srsx-3* in both AWC neurons, leading to the 2AWC^OFF^ phenotype (Figures 1Aiii, 1B, rows b-d and o, and S1A). The *ajm-1(vy11)* allele is temperature-sensitive for the 2AWC^OFF^ phenotype, with 32% of the animals showing 2AWC^OFF^ at 15℃ (Figure 1B, row c), 75% at 20℃ (Figure 1B, row b), and 92% at 25℃ (Figure 1B, row d).

**Figure 1.**
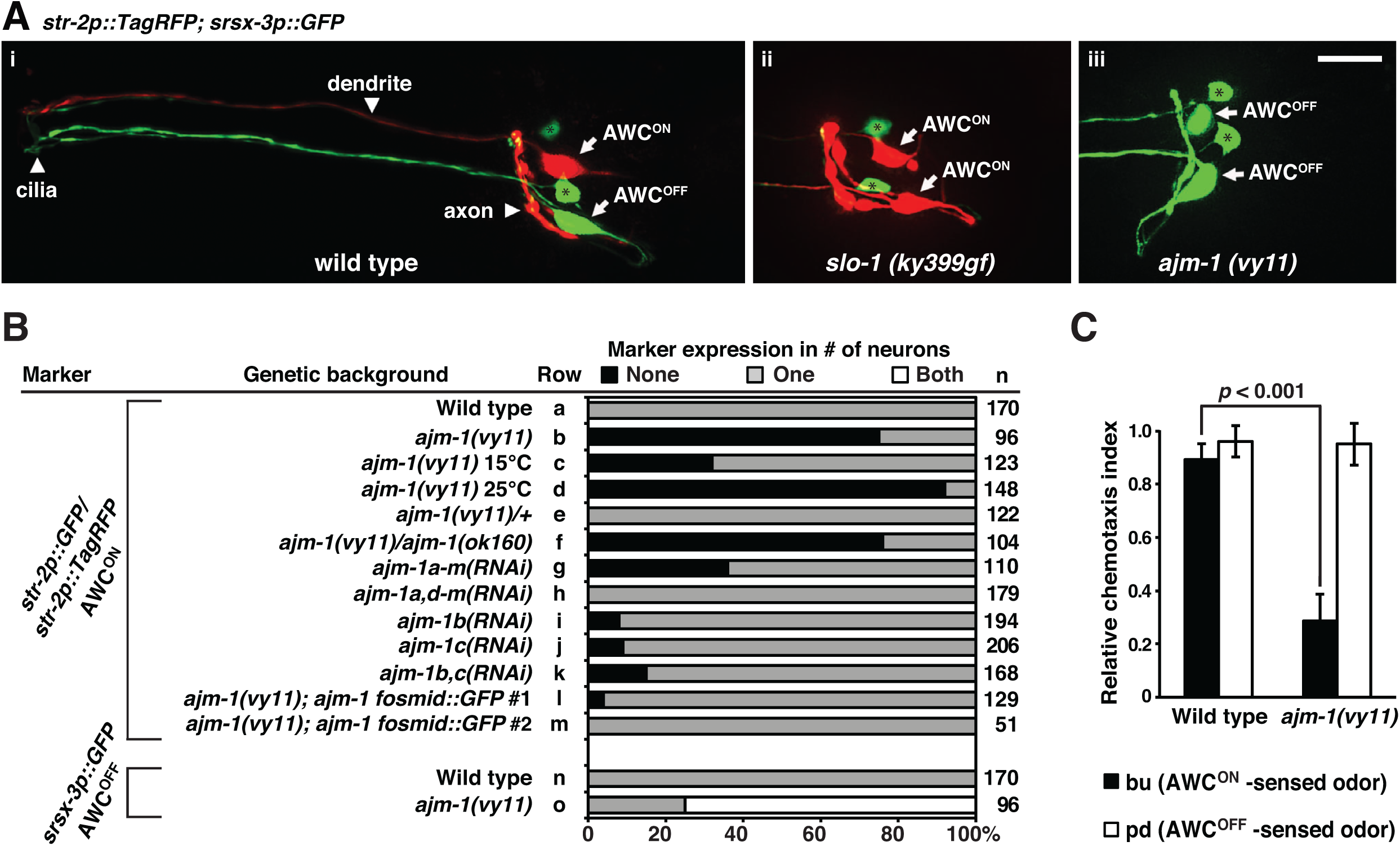
*ajm-1* is required to promote the AWC^ON^ subtype. **(A)** Images of wild type (i), *slo-1(ky399gf)* mutants (ii), and *ajm-1(vy11)* mutants (iii) expressing the transgene *str-2p::TagRFP* (AWC^ON^ marker); *srsx-3p::GFP* (AWC^OFF^ marker) in the adult stage. (i) Wild-type animals express *str-2p::TagRFP* in the AWC^ON^ cell and *srsx-3p::GFP* in the AWC^OFF^ cell. (ii) *slo-1(ky399gf)* mutants lose expression of *srsx-3p::GFP* and express *str-2p::TagRFP* in both AWC neurons, resulting in a 2AWC^ON^ phenotype. (iii) *ajm-1(vy11)* mutants lose expression of *str-2p::TagRFP* and express *srsx-3p::GFP* in both AWC neurons, resulting in a 2AWC^OFF^ phenotype. *srsx-3p::GFP* is also expressed in 2 AWB neurons. Arrows indicate AWC cell bodies; asterisks indicate AWB cell bodies. Anterior is left, and ventral is down. Scale bar, 10 μm. **(B)** Expression of AWC^ON^ and AWC^OFF^ markers in adults. RNA interference (RNAi) targeting different *ajm-1* isoforms was performed in the RNAi-sensitive strain *eri-1(mg366); lin-15B(n744)* double mutant, as described (Kamath, Martinez-Campos et al. 2001, Sieburth, Ch’ng et al. 2005). n, total number of animals scored. Animals were grown at 20℃, unless otherwise indicated. **(C)** Quantification of the chemotaxis index of wild type and *ajm-1(vy11)* mutants to AWC^ON^-and AWC^OFF^-sensed odors. bu, butanone; pd, 2,3-pentanedione. A higher chemotaxis index indicates a higher level of attraction. Student’s *t*-test was used to determine the *p*-value. Error bars represent the standard error of the mean (SEM).

To examine the general AWC neuron identity in the *vy11* mutants, the expression of two general AWC identity markers, the guanylyl cyclase *odr-1* and the homeodomain transcription factor *ceh-36*, was analyzed. The *odr-1* and *ceh-36* markers were expressed in both AWC neurons in wild-type animals and *vy11* mutants (Figure S1B). These results suggest that the *vy11* mutation does not affect general AWC cell identity but does cause a defective AWC asymmetry phenotype.

The AWC subtypes respond to different odors: the AWC^ON^ neuron detects butanone, while the AWC^OFF^ neuron detects 2,3-pentanedione (Wes and Bargmann 2001). Chemotaxis assays were performed to determine whether *vy11* mutants are defective in sensing the AWC^ON^ odorant butanone. Wild-type animals were attracted to both the AWC^ON^- and AWC^OFF^-sensed odors (Figure 1C). *vy11* mutants were attracted normally to the AWC^OFF^-sensed odor 2,3-pentanedione, but their ability to chemotaxis toward the AWC^ON^-sensed odor butanone was significantly reduced (*p* < 0.001) (Figure 1C). Taken together, these results suggest that the *vy11* mutation leads to a loss of the AWC^ON^ neuron subtype at both the molecular and functional levels.

### *vy11* is a missense mutation in *ajm-1* (apical junction molecule 1)

We identified the molecular lesion in *vy11* mutants using one-step whole genome sequencing and single-nucleotide polymorphism (SNP) mapping (Doitsidou, Poole et al. 2010). The *vy11* lesion is a C to T mutation, resulting in a glutamine to ochre stop change in the first exon of C25A11.4a/*ajm-1a* isoform (Figures 2A and S2A). *ajm-1* has 13 alternatively-spliced isoforms (wormbase.org) (Figures 2A and S2A). The glutamine residue affected by the *vy11* mutation is present in *ajm-1a-d, f, g, k-m*, but absent in *ajm-1e, h, i, j*.

**Figure 2.**
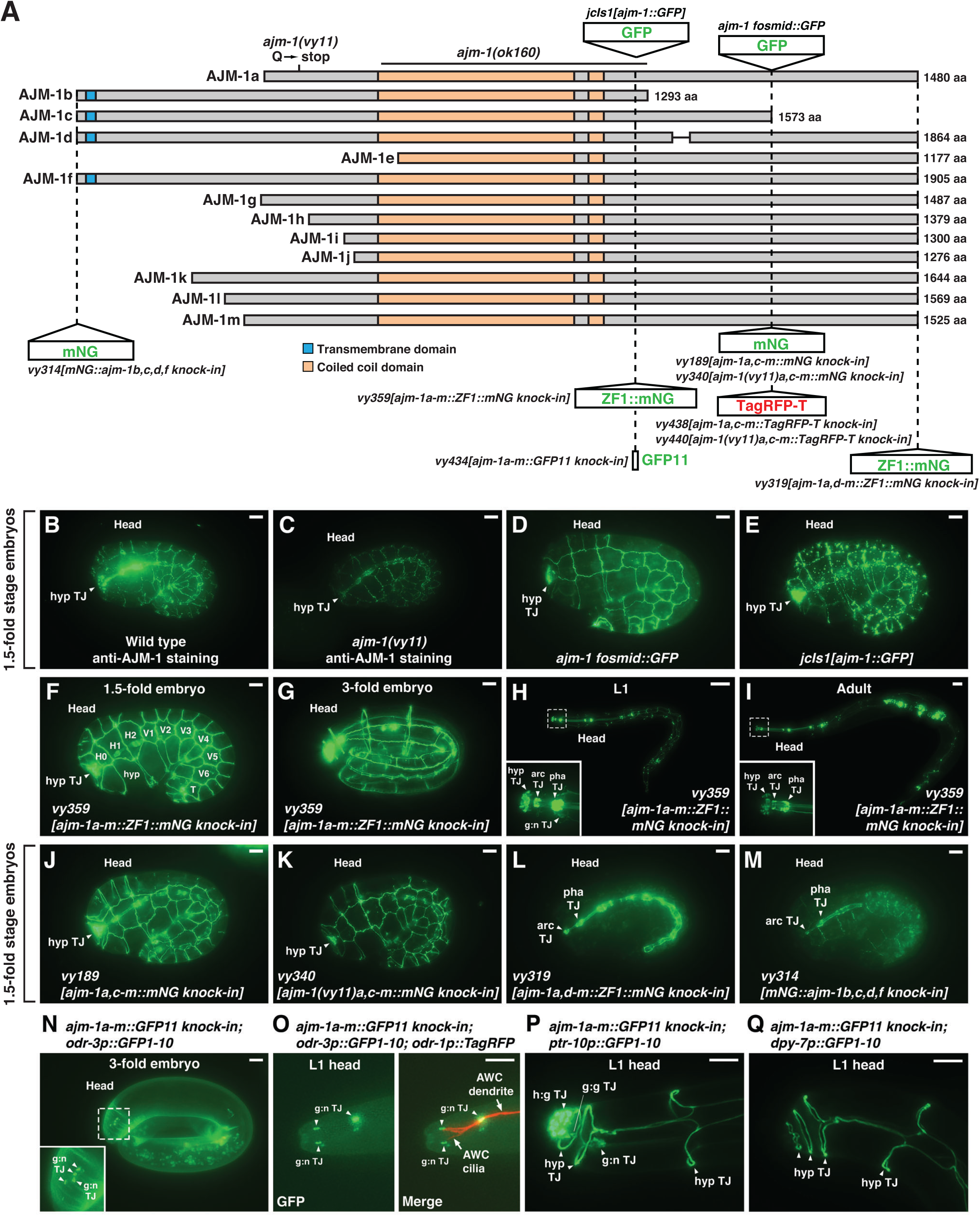
AJM-1 is broadly expressed in multiple tissues. (**A**) Structure of *C. elegans* AJM-1 protein isoforms, a-m. The length of each isoform is indicated by the number of amino acids (Ackley, Crew et al.). The *ajm-1* fosmid::GFP includes a 1063 bp upstream region, the *ajm-1a* isoform coding region with GFP inserted after exon 13, and a 15 kb downstream region. Dashed lines indicate the location of GFP-fusion transgenes or CRISPR/Cas9 knock-in tags. ZF1, PIE-1 C-C-C-H type zinc-finger domain. *mNG*, mNeonGreen. (**B, C**) Images of wild-type (B) and *ajm-1(vy11)* (C) 1.5-fold stage embryos stained with anti-AJM-1 antibodies. (**D**, **E**) Images of 1.5-fold stage embryos expressing *ajm-1 fosmid::GFP* (D) and *ajm-1::GFP* (E) transgenes. (**F-I**) Images of a 1.5-fold stage embryo (F), a 3-fold stage embryo (G), a first-stage larva (L1) (H), and an adult (I) expressing *ZF1::mNG knock-in* in all AJM-1 isoforms a-m. (**J**, **K**) Images of wild-type (J) and *ajm-1(vy11)* (K) 1.5-fold stage embryos expressing *mNG knock-in* in AJM-1 isoforms a,c-m. AJM-1b isoform is not tagged. The hypodermal expression of mNG remains strong in wild type (J) when compared to (F). (**L**) Image of a 1.5-fold stage embryo expressing *ZF1::mNG knock-in* at the C-terminus of AJM-1 isoforms a,d-m. AJM-1 isoforms b and c are not tagged. The expression of mNG is not detected in hypodermal cells when compared to (F) and (J). (**M**) Image of a 1.5-fold stage embryo expressing *mNG knock-in* tagged at the N-terminus of AJM-1 isoforms b, c, d, and f. Other AJM-1 isoforms are not tagged. The expression of mNG is still detected in hypodermal cells but appears dimmer than in (F) and (J). (**N**-**Q**) Tissue-specific visualization of endogenous AJM-1 protein expression using split GFP in specific tissues. GFP11 is knocked in all AJM-1 isoforms a-m. GFP1-10 is under the control of a tissue-specific promoter expressed in AWC neurons (N and O), pan-glial cells (P), or hypodermal cells (Q). **(B-Q)** hyp TJ, hypodermal tight junction; H0-H2, V1-V6, and T, embryonic seam cells; hyp, hypodermal cell; arc TJ, tight junction in arcade cells; pha TJ, pharyngeal tight junction; g:n TJ, tight junction between glial cells and neurons; h:g TJ, tight junction between hypodermal and glial cells; g:g TJ, glial tight junction. Scale bars, 5 μm (B-G, J-Q) and 20 μm (H, I).

*ajm-1* encodes an apical junction molecule, an ortholog of human AJM1 (apical junction component 1) (wormbase.org). It contains a coiled-coil domain that is shared by all isoforms. AJM-1b, c, d, and f isoforms are predicted to have one transmembrane domain region starting at amino acid position 21 and ending at position 43 using the Simple Modular Architecture Research Tool (SMART) (Letunic, Doerks et al. 2015) (Figure 2A). AJM-1 is important for the integrity of the apical junctional domain of the *C. elegans* epithelia (Koppen, Simske et al. 2001, Lockwood, Lynch et al. 2008).

The *ajm-1(ok160)* mutation results in a deletion of almost the entire coiled-coil domain of all isoforms (Figure 2A). *ajm-1(ok160)* mutants cannot survive beyond the 2-fold embryonic stage (Koppen, Simske et al. 2001), obstructing AWC phenotype scoring in larvae. However, *ajm-1(vy11)*, which affects 9 of 13 isoforms, is viable, suggesting that *ajm-1(vy11)* is likely not a null allele. Transheterozygotes of *vy11* and *ajm-1(ok160)* null alleles showed a similar percentage of the 2AWC^OFF^ phenotype as the *vy11* homozygous mutants (Figure 1B, rows b and f). Heterozygous *ajm-1(vy11/+)* animals displayed wild-type AWC asymmetry, indicating that the *ajm-1(vy11)* allele is recessive (Figure 1B, row e). Together, these results suggest that *vy11* is a reduction-of-function allele.

RNA interference (RNAi) targeting all *ajm-1* isoforms caused embryonic lethality in most animals. However, the 2AWC^OFF^ phenotype was observed in rare animals that escaped lethality caused by *ajm-1a-m(RNAi)* (Figure 1B, row g). Similar results were observed in RNA interference targeting *ajm-1b* (Figure 1B, row i)*, ajm-1c* (Figure 1B, row j), or both isoforms together (Figure 1B, row k). However, RNAi targeting other isoforms, except b and c, resulted in a wild-type AWC asymmetry phenotype (Figure 1B, row h). Taken together, these results suggest that *ajm-1b* and *ajm-1c* isoforms are the major isoforms required for AWC asymmetry.

An *ajm-1* fosmid::GFP clone almost completely rescued the 2AWC^OFF^ phenotype in *vy11* mutants (Figures 1B, rows l and m, and 2A), further supporting that the identified mutation in *ajm-1* causes the *vy11* 2AWC^OFF^ phenotype.

### AJM-1 is broadly expressed in multiple tissues

In a previous study, antibody staining and an integrated *juIs1[ajm-1::GFP]* transgene were used to demonstrate AJM-1 localization to the apical domain of epithelial junctions. This localization was shown in the embryonic hypodermis, pharynx, and intestine, as well as in post-embryonic epithelia (Koppen, Simske et al. 2001). In addition, AJM-1 has been shown to be localized at the three distinct electron-dense tight junctions between amphid neurons and sheath glia, between sheath and socket glia, and between socket glia and hypodermal cells at the anterior tip of the animal (Nguyen, Liou et al. 2014, Nechipurenko, Olivier-Mason et al. 2016, Low, Williams et al. 2019).

Consistent with previous findings, we observed a similar expression pattern of AJM-1 in glial cells, the hypodermis, the pharynx, and the intestine, as revealed by antibody staining in wild-type 1.5-fold stage embryos (Figure 2B). Anti-AJM-1 staining in *ajm-1(vy11)* 1.5-fold stage embryos showed a similar pattern but with a significantly reduced expression level (Figure 2C). These results suggest that the *vy11* mutation reduces the expression level and/or the stability of AJM-1. The expression pattern of AJM-1 in glial cells, hypodermis, pharynx, and intestine was also observed in an extrachromosomal *ajm-1 fosmid::GFP* reporter strain (Figure 2D) and confirmed in the integrated *juIs1[ajm-1::GFP]* strain (Figure 2E).

To further determine the endogenous expression pattern of all *ajm-1* isoforms, we generated *ajm-1a-m::ZF1::mNG knock-in* (where [mNG] represents mNeonGreen) using Cas9-triggered homologous recombination (Dickinson, Ward et al. 2013, Armenti, Lohmer et al. 2014, Dickinson, Pani et al. 2015, Dickinson and Goldstein 2016) (Figure 2A). *ajm-1a-m::ZF1::mNG knock-in* animals showed a comparable expression pattern to AJM-1 antibody staining at the 1.5-fold (Figure 2B and 2F), with continuous expression in glial cells, hypodermis, pharynx, and intestine from the 3-fold stage (Figure 2G) through the first larval stage (Figure 2H) to the adult stage (Figure 2I). We also generated *ajm-1a,c-m::mNG knock-in*, which tagged 12 out of 13 AJM-1 isoforms, omitting AJM-1b (Figure 2A and 2J). There was no obvious difference in the expression pattern of *ajm-1a,c-m::mNG knock-in* in comparison to *ajm-1a-m::ZF1::mNG knock-in* at the 1.5-fold stage (Figure 2F and 2J). In *ajm-1(vy11)* mutants, *ajm-1(vy11)a,c-m::mNG knock-in* showed a similar expression pattern compared to the wild type but with decreased mNG fluorescence intensity (Figure 2J and 2K). *ajm-1a,d-m::ZF1::mNG knock-in* tagged the C-terminal end of 11 endogenous AJM-1 isoforms, omitting AJM-1b and c (Figure 2A). *ajm-1a,d-m::ZF1::mNG knock-in* animals showed lost mNG expression in glial and hypodermal cells (Figure 2L), compared to *ajm-1a-m::ZF1::mNG knock-in* and *ajm-1a,c-m::mNG knock-in*. These results suggest that *ajm-1b* and *ajm-1c*, the major isoforms required for AWC asymmetry (Figure 1B, rows g-k), are the primary isoforms expressed in glial and hypodermal cells. *mNG::ajm-1b,c,d,f knock-in,* which tagged endogenous AJM-1b, c, d, and f isoforms at the N-terminal end (Figure 2A), showed mNG expression in the intestine and hypodermis with reduced intensity (Figure 2M). All aforementioned *ajm-1::mNG knock-ins* are functional in AWC asymmetry, as the *knock-ins* displayed a wild-type 1AWC^ON^/1AWC^OFF^ phenotype (Figure S2B, rows c-f).

To facilitate dissecting the tissues expressing AJM-1 and its subcellular localization within those tissues, we used a tissue-specific split-GFP approach (Cabantous, Terwilliger et al. 2005, Noma, Goncharov et al. 2017, Goudeau, Sharp et al. 2021). We generated the *ajm-1a-m::GFP11 knock-in*, tagging all *ajm-1* isoforms using CRISPR/Cas9-triggered homologous recombination and single-stranded oligodeoxynucleotide donors (ssODN) (Farboud, Severson et al. 2019, Goudeau, Sharp et al. 2021) (Figure 2A). *ajm-1a-m::GFP11* is functional in AWC asymmetry (Figure S2B, row h). GFP1-10 was expressed from tissue-specific promoters and crossed with *ajm-1a-m::GFP11 knock-in* strains. When GFP1-10 was expressed from the AWC-specific *odr-3* promoter (Roayaie, Crump et al. 1998), reconstituted AJM-1a-m::GFP was expressed at the tight junctions between glial cells and neurons near the anterior of the head in a 3-fold stage embryo (Figure 2N). To further examine the subcellular localization of AJM-1 in AWC, the *odr-1p::TagRFP* transgene (Cochella, Tursun et al. 2014) was used to label the entirety of both AWC neurons. In the first-stage larvae, reconstituted AJM-1a-m::GFP was expressed at the tight junctions between glial cells and neurons near the AWC dendrite endings and the tip of the AWC cilia (Figure 2O). When GFP1-10 was expressed from the glial cell-specific *ptr-10* promoter (Yoshimura, Murray et al. 2008), reconstituted AJM-1a-m::GFP expression was detected at hypodermal tight junctions, the tight junction between hypodermal and glial cells, the glial tight junction, and the tight junction between glial cells and neurons in the head of first-stage larvae and 3-fold embryos (Figures 2P and S2C). Similar expression of reconstituted AJM-1a-m::GFP at the hypodermal tight junctions was observed when GFP1-10 was expressed from the hypodermal cell-specific *dpy-7* promoter (Gilleard, Barry et al. 1997) (Figures 2Q and S2D). Together, these results indicate that AJM-1 is broadly expressed at the apical junctions of multiple tissues, including AWC neurons, glial cells, and hypodermal cells.

### The localization of AJM-1 at the apical region of cell junctions is reduced in *ajm-1(vy11)* mutants

The correct localization of AJM-1 at the apical region of the lateral cell-cell junction is essential for its function (Koppen, Simske et al. 2001, McMahon, Legouis et al. 2001, Lockwood, Lynch et al. 2008). To determine whether the *ajm-1(vy11)* mutation affects the proper subcellular distribution of AJM-1 protein at the apical junction, we generated the *ajm-1a,c-m::TagRFP-T knock-in* and *ajm-1(vy11)a,c-m::TagRFP-T knock-in* using Cas9-triggered homologous recombination (Dickinson, Ward et al. 2013, Dickinson, Pani et al. 2015, Dickinson and Goldstein 2016) (Figure 2A). Additionally, the β-G spectrin reporter, *GFP::unc-70 knock-in*, was used to mark the basal extent of the lateral cell membrane in 1.5-fold embryos (Moorthy, Chen et al. 2000, McMahon, Legouis et al. 2001, Jia, Li et al. 2019). To better differentiate individual cells in 1.5-fold embryos, a single-copy transgene *oxTi973[eft-3p::GFP::2xNLS::tbb-2 3’UTR]* (*Caenorhabditis* Genetics Center) was used to mark the nucleus. Finally, to analyze the position of the AJM-1 protein in the lateral membrane, a single-pixel transverse view was generated from the cross-section of one hypodermal cell in both the wild-type and *ajm-1(vy11)* 1.5-fold embryos (Figure 3A & 3C). AJM-1 protein was appropriately located at the apical region of the lateral membrane in both wild-type animals and *ajm-1(vy11)* mutants. However, the expression level of AJM-1 was reduced in *ajm-1(vy11)* mutants (Figure 3A’, 3A”, 3C’, and 3C”). Together, these results suggest that the *ajm-1(vy11)* mutation reduces the subcellular distribution of AJM-1 protein at the apical junction.

**Figure 3.**
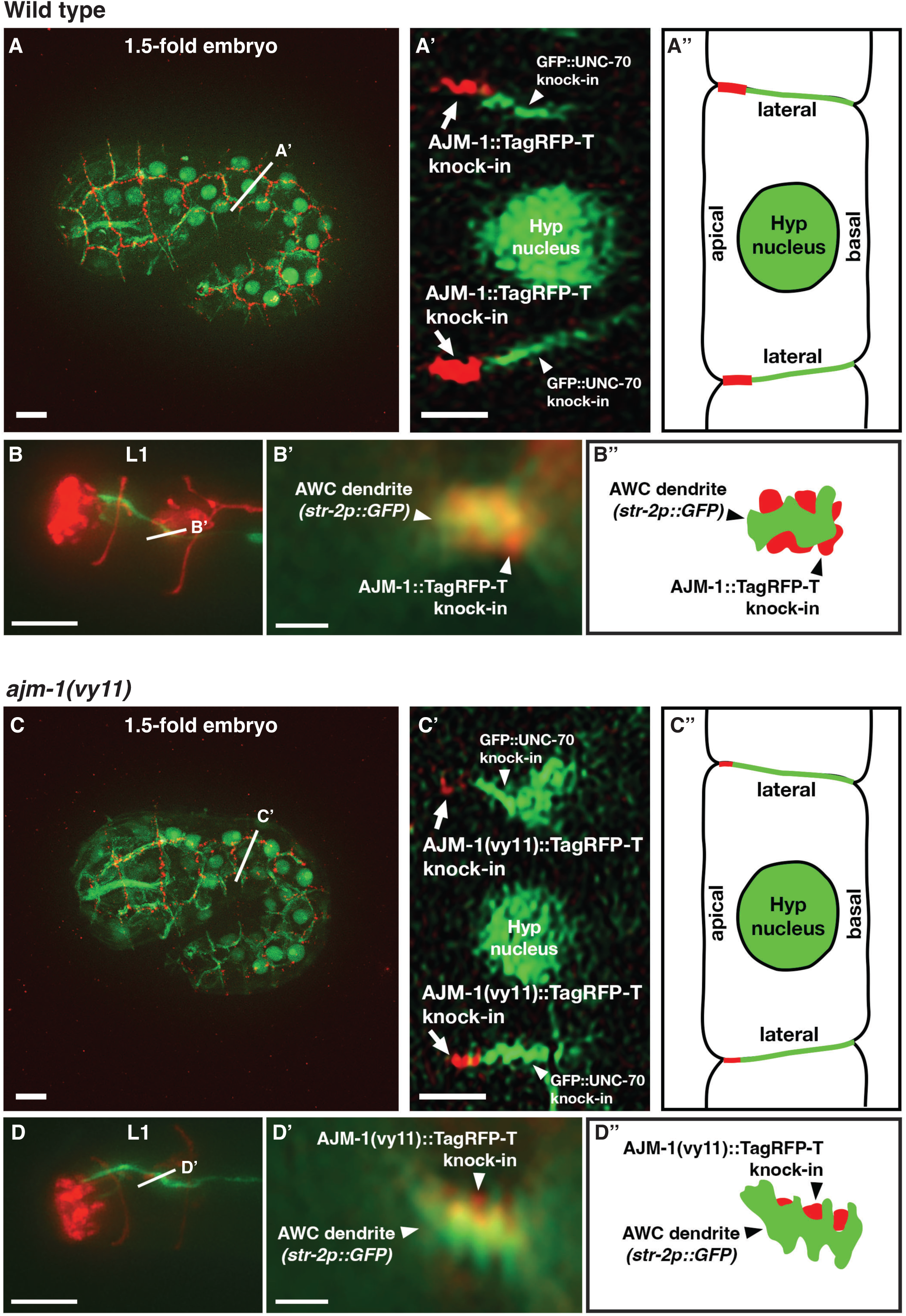
The localization of AJM-1 at the apical region of cell junctions is reduced in *ajm-1(vy11)* mutants. **(A**, **C)** Images of wild-type (A) and *ajm-1(vy11)* (C) 1.5-fold embryos expressing *ajm-1a,c-m::TagRFP-T knock-in*, *unc-70(cas962[GFP::unc-70 knock-in])* in the cell membrane, and *oxTi973[eft-3p::GFP::2xNLS::tbb-2 3’UTR + HygroR]* in the nucleus. Straight white lines indicate the position of transverse sections of a hypodermal cell. Scale bars, 5 μm. **(A’, C’)** Transverse views of hypodermal cells showing the localization of AJM-1::TagRFP-T at the apical region of the lateral cell membrane in wild type (A’) and *ajm-1(vy11)* (C’). The expression level of AJM-1::TagRFP-T was decreased in the *ajm-1(vy11)* embryo. Scale bars, 2 μm. **(A”, C”)** Schematic diagram of (A’) and (C’). **(B, D)** Images of wild-type (B) and *ajm-1(vy11)* (D) first-stage larvae (L1) expressing *ajm-1a,c-m::TagRFP-T knock-in* and *str-2p::GFP* (labeling the AWC^ON^ dendrite and cilia. White straight lines indicate the position of transverse sections of the AWC dendrite. Scale bars, 5 μm. **(B’, D’)** Transverse views showing the localization of AJM-1::TagRFP-T adjacent to the AWC dendrite in wild type (B’) and *ajm-1(vy11)* (D’). The expression level of AJM-1::TagRFP-T was decreased in the *ajm-1(vy11)* L1. Scale bars, 1 μm. **(B”, D”)** Schematic diagram of (B’) and (D’). **(A’-D’)** Z-axis rotations of a single transverse slice through the section indicated by the straight white line in images A-D.

Since our tissue-specific visualization of AJM-1 indicated the protein was expressed near the dendrite endings of AWC neurons in first-stage larvae (Figure 2O), we determined whether this pattern was affected by the *ajm-1(vy11)* mutation. Similar transverse images were acquired at the overlapping area between AJM-1::TagRFP-T and the AWC dendrite labeled with a GFP marker (Figure 3B and 3D). The AJM-1 localization was at a similar position near the AWC dendrite endings in both wild-type and *ajm-1(vy11)* larvae, but at a reduced level in the mutants (Figure 3B’, 3B”, 3D’, and 3D”).

### *ajm-1* functions in neuronal and non-neuronal tissues to promote AWC^ON^

To determine the tissues required for *ajm-1* function in AWC asymmetry, we performed rescue experiments by expressing *ajm-1* cDNA from different tissue-specific promoters in *ajm-1(vy11)* mutants (Figure 4). The *odr-3p::ajm-1* transgene driven by the AWC-specific *odr-3* promoter (Roayaie, Crump et al. 1998) was not sufficient to fully rescue the *ajm-1(vy11)* 2AWC^OFF^ mutant phenotype (Figure 4, data group c, compared to data groups a and b). This result suggests that *ajm-1* may also act in non-AWC cells to regulate AWC asymmetry. Similar rescue results were observed in the *rab-3p::ajm-1* transgene driven by the pan-neuronal *rab-3* promoter (Stefanakis, Carrera et al. 2015) (Figure 4, data group d, compared to data group c). Taken together, these results suggest that *ajm-1* may also act in non-neuronal cells to promote the AWC^ON^ subtype.

**Figure 4.**
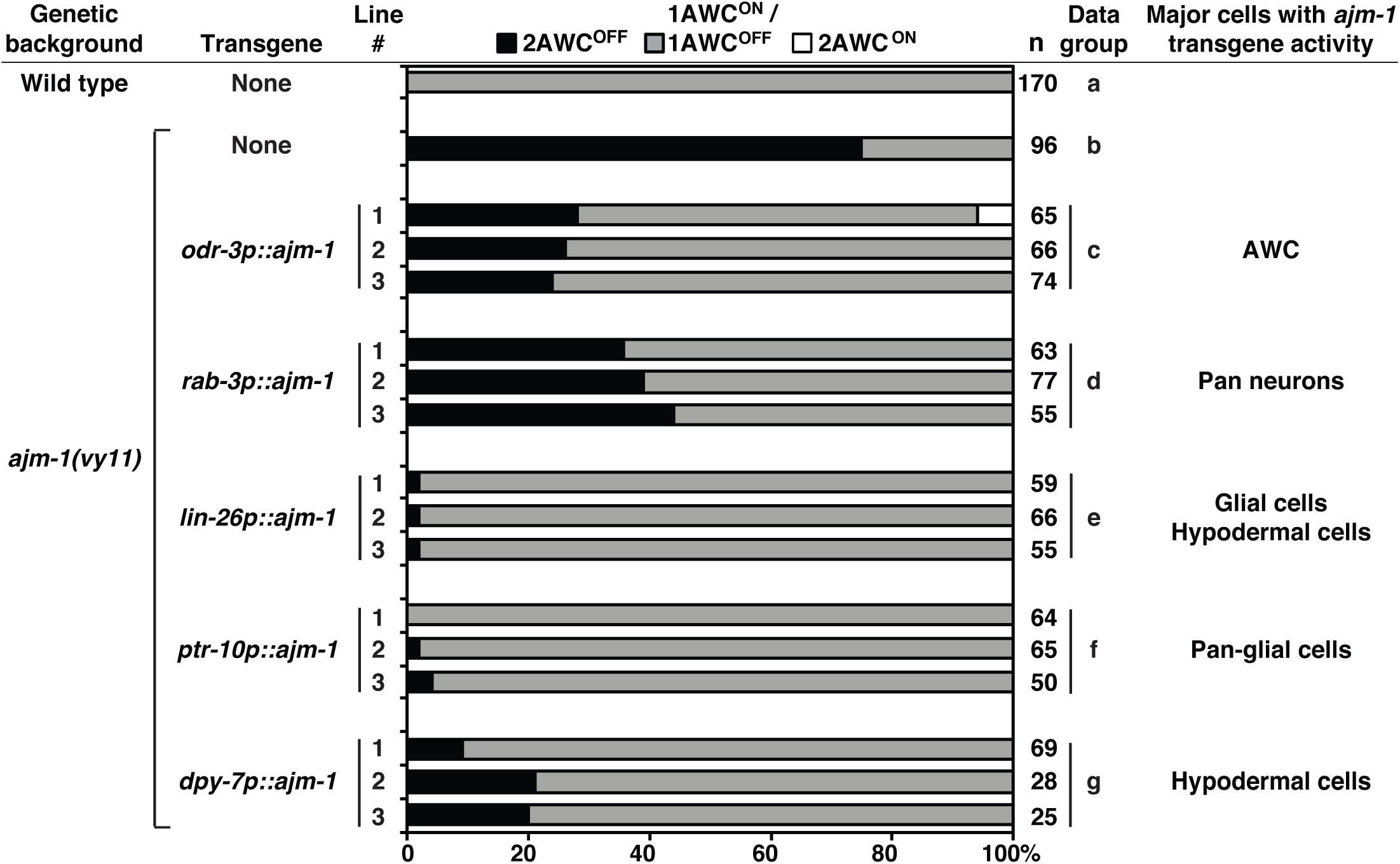
*ajm-1* functions in neuronal and non-neuronal tissues to promote AWC^ON^. Quantification of AWC asymmetry phenotypes in three independent transgenic lines expressing *ajm-1a e1-13::SL2::myrGFP*, which consists of the *ajm-1a* cDNA region spanning from the first ATG of *ajm-1a* exon 1 to the first 144 bp of *ajm-1a* exon 13 and a SL2 trans-splicing sequence followed by myrGFP, from different tissue-specific promoters in *ajm-1(vy11)* mutants. Animals were scored at the adult stage. n, total number of animals scored. The AWC phenotype data of the wild type (row a) and *ajm-1(vy11)* (row b) are the same as those of Figure 1B (rows a and b, respectively).

Since AJM-1 was broadly detected in glial and hypodermal cells (Figures 2B-2Q, S2C, and S2D), the *lin-26p::ajm-1* transgene driven by a glial and hypodermal cell-specific *lin-26* promoter (Labouesse, Sookhareea et al. 1994, Labouesse, Hartwieg et al. 1996) was used for rescue experiments. The *lin-26p::ajm-1* transgene almost completely rescued the *ajm-1(vy11)* 2AWC^OFF^ mutant phenotype (Figure 4, data group e), suggesting that *ajm-1* may function in glial and/or hypodermal cells to promote AWC^ON^.

We then determined whether expression of *ajm-1* in glial or hypodermal cells alone was sufficient to rescue the *ajm-1(vy11)* 2AWC^OFF^ mutant phenotype. The *ptr-10::ajm-1* transgene driven by a glial cell-specific *ptr-10* promoter (Yoshimura, Murray et al. 2008) rescued the *ajm-1(vy11)* 2AWC^OFF^ mutant phenotype to a level similar to *lin-26p::ajm-1* (Figure 4, data group f, compared to data group e). This result suggests that the *ajm-1* function in glial cells is sufficient for promoting AWC^ON^. On the other hand, the *dpy-7p::ajm-1* transgene driven by a hypodermal cell-specific *dpy-7* promoter (Gilleard, Barry et al. 1997) had a lower rescue ability than *lin-26p::ajm-1* and *ptr-10p::ajm-1* (Figure 4, data group g, compared to data groups e and f). This result suggests that *ajm-1* function in hypodermal cells alone may not be sufficient for promoting AWC^ON^.

Taken together, these results suggest that *ajm-1*, in addition to its cell-autonomous role in AWC cells, has a non-cell-autonomous role in non-neuronal tissues, including glial and hypodermal cells, to promote the AWC^ON^ subtype.

### *ajm-1* acts non-cell autonomously to promote AWC^ON^

To further support the non-cell-autonomous role of *ajm-1* in AWC asymmetry, we performed genetic mosaic analysis on *ajm-1(vy11)* mutant animals expressing one of the three rescue transgenes from extrachromosomal arrays, *ajm-1 fosmid::GFP*, *lin-26p::ajm-1*, or *ptr-10p::ajm-1* (Figure 5A). Mosaic animals were generated by a spontaneous and random loss of unstable extrachromosomal transgenes during mitosis. The *odr-1p::DsRed* reporter transgene (expressed in two AWC and two AWB neurons), co-injected with the rescue transgene, was used as a mosaic marker. The pairs of AWC and AWB neurons are derived from 4 different cell lineages, each of which differentiates into various cell types during development, including neurons, glial cells, hypodermal cells, excretory cells, and muscle cells (Figure 5B). The expression of *odr-1p::DsRed* in one of the four cells was used to infer the presence of the co-injected rescue transgene in other cell types of the same cell lineage. We specifically selected mosaic animals that retained the transgene in only a single cell lineage, as indicated by *odr-1p::DsRed* expression in AWCL, AWCR, AWBL, or AWBR, but lost the transgene from the other three lineages.

**Figure 5.**
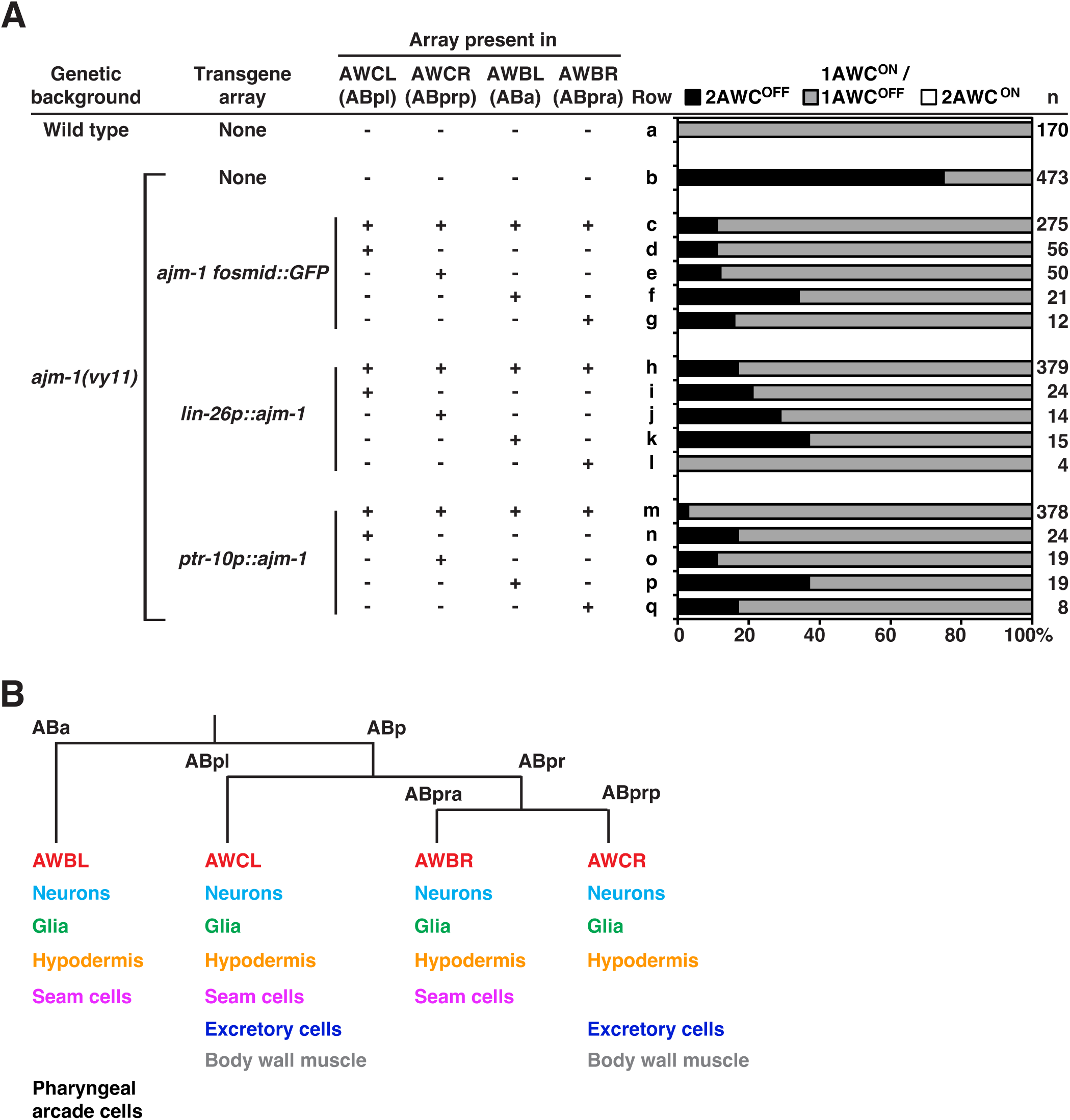
*ajm-1* acts non-cell autonomously to promote AWC^ON^. **(A)** Quantification of AWC asymmetry phenotypes in *ajm-1(vy11)* mosaic animals containing the extrachromosomal transgene array *ajm-1 fosmid::GFP*, *lin-26p::ajm-1* (expressing *ajm-1f* cDNA from the *lin-26* promoter), or *ptr-10p::ajm-*1 (expressing *ajm-1a e1-13::SL2::myrGFP* from the *ptr-10* promoter) in cells derived from different cell lineages, inferred by the presence of co-injected *odr-1p::DsRed* marker (expressed in 4 cells representing different cell lineages). AWCL, left AWC; AWCR, right AWC; AWBL, left AWB; AWBR, right AWB. +, presence of extrachromosomal array. -, absence of extrachromosomal array. Animals were scored at the adult stage. n, total number of animals scored. The AWC phenotype data of the wild type (row a) are the same as those of Figure 1B (row a). **(B)** Simplified cell lineage of *C. elegans* starting at the second cell division, with derived tissue types. For mosaic analysis, individual cell lineages were traced by *odr-1p::DsRed* expression in AWBL, AWBR, AWCL, or AWCR (labeled in red). Other neurons are indicated in cyan.

The presence of *ajm-1 fosmid::GFP* in all AWC and AWB lineages rescued the *ajm-1(vy11)* 2AWC^OFF^ mutant phenotype from 75% to 11% penetrance (Figure 5A, row c, compared to rows a and b). Transgene arrays retained only in the AWCL (ABpl) (Figure 5A, row d), AWCR (ABprp) (Figure 5A, row e), AWBL (ABa) (Figure 5A, row f), or AWBR lineage (ABpra) (Figure 5A, row g) also rescued the *ajm-1(vy11)* 2AWC^OFF^ mutant phenotype to 11%-34% penetrance. Similar mosaic analysis results were observed with the *lin-26p::ajm-1* and *ptr-10p::ajm-1* rescue transgenes. The *ajm-1(vy11)* 2AWC^OFF^ mutant phenotype was rescued by the presence of the *lin-26p::ajm-1* or the *ptr-10p::ajm-1* transgene in all four AWC and AWB lineages (Figure 5A, rows h and m) to 17% or 3% penetrance, respectively. Retention of the transgene array in any one of the four cell lineages alone (Figure 5A, rows i-l and n-q) also rescued the *ajm-1(vy11)* 2AWC^OFF^ mutant phenotype to 0%-37% penetrance. Taken together, these results suggest that *ajm-1* function in each of the four cell lineages may be redundant for AWC asymmetry. These results, consistent with the rescue data (Figure 4), also support the non-cell-autonomous function of *ajm-1* in non-AWC cells in promoting AWC^ON^.

### *ajm-1* acts embryonically in AWC asymmetry

To determine the temporal requirement of *ajm-1* in AWC asymmetry, we used the ZF1-mediated degradation system (Armenti, Lohmer et al. 2014) to degrade endogenous AJM-1 protein at different developmental stages from embryogenesis to larvae. We generated the *ajm-1a-m::ZF1::mNG knock-in* strain, in which all *ajm-1* isoforms were tagged with the 36 amino acid PIE-1 C-C-C-H type zinc-finger domain ZF1 (Reese, Dunn et al. 2000) fused with mNG fluorescent protein (Figures 2A and S2A) using CRISPR/Cas9 genome editing (Dickinson, Ward et al. 2013, Dickinson, Pani et al. 2015, Dickinson and Goldstein 2016).

Heterologous cytosolic or transmembrane proteins fused with a ZF1 domain can bind to the E3 ubiquitin ligase substrate-recognition subunit ZIF-1 and subsequently be recruited to an ECS (Elongin-C, Cul2, SOCS-box family) E3 ubiquitin ligase complex for proteosome-mediated degradation (Armenti, Lohmer et al. 2014). The *ajm-1(ok160)* null allele causes embryonic lethality (Koppen, Simske et al. 2001). Therefore, constitutive degradation of AJM-1 may cause animal lethality and prevent the quantification of AWC asymmetry phenotypes. To bypass this issue, we used the heat-shock responsive *hsp-16.2* promoter (Bacaj and Shaham 2007) to drive *zif-1* expression, thereby degrading AJM-1::ZF1::mNG upon heat-shock treatment at specific developmental stages. The effect of AJM-1 degradation, which was indicated by the loss of AJM-1::ZF1::mNG protein expression, on AWC asymmetry was analyzed.

After heat-shock treatment, *ajm-1a-m::ZF1::mNG knock-in* embryos containing the *hsp-16.2p::zif-1* extrachromosomal transgene showed obvious mNG degradation, whereas non-transgenic embryos showed no degradation of mNG expression (Figure S3A and S3B). In addition, heat shock of mixed-stage embryos of transgenic lines resulted in 57%-63% penetrance of the 2AWC^OFF^ phenotype (Figure 6A, data group c), compared to 75% in *ajm-1(vy11)* mutants (Figure 6A, data group b). In contrast, control experiments showed that non-transgenic embryos with or without heat shock and transgenic lines without heat shock showed no or very low (<1%) incidence of the 2AWC^OFF^ phenotype (Figure S4A, rows c-e).

**Figure 6.**
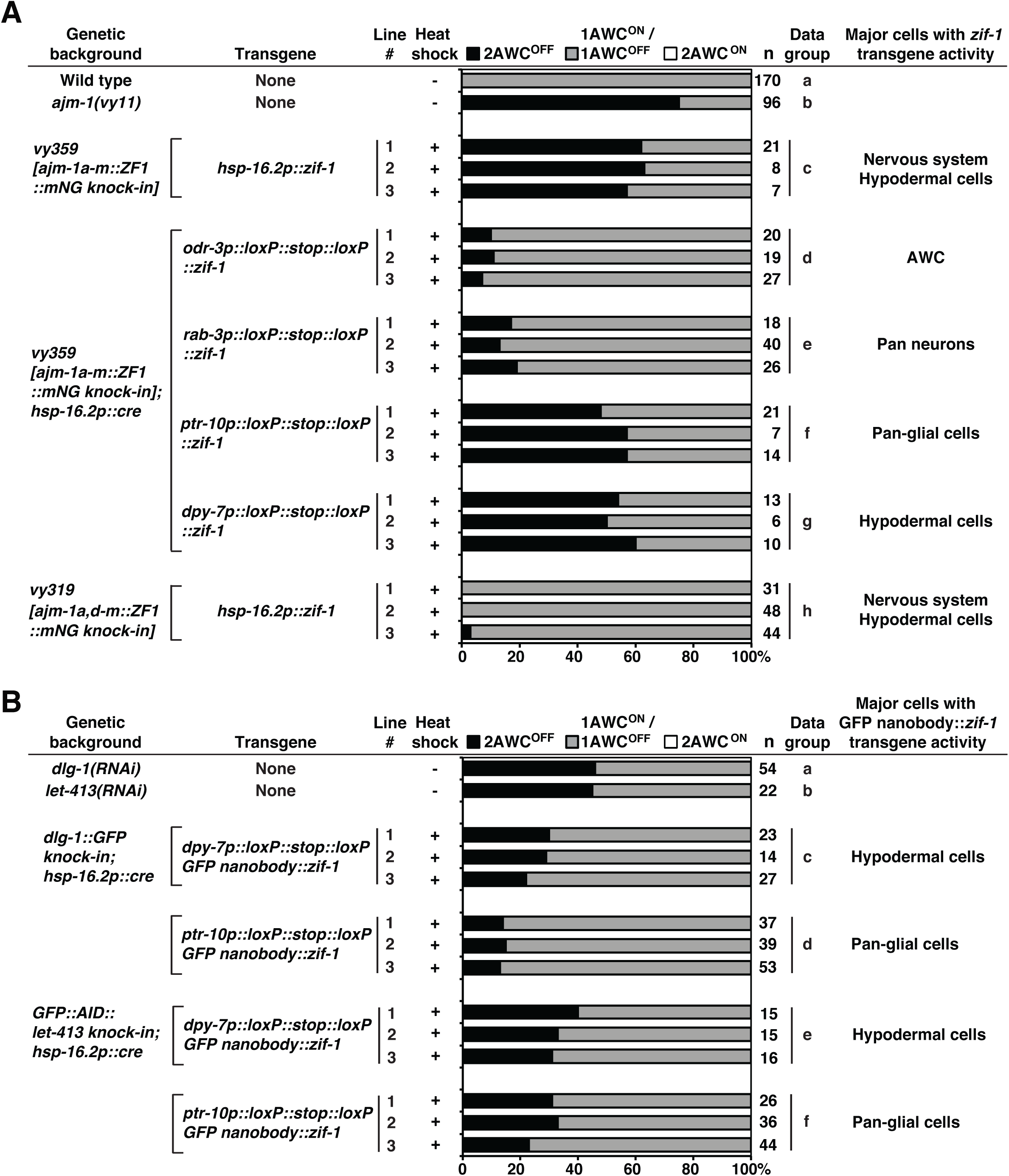
*ajm-1* is required in hypodermal and glial cells to promote AWC^ON^. **(A)** Quantification of AWC asymmetry phenotypes in animals expressing *ZF1::mNG* knock-in in all AJM-1 isoforms a-m or at the C-terminus of AJM-1 isoforms a,d-m, with transgenes of ZIF-1 (the substrate-binding subunit of the E3 ligase) driven by different tissue-specific promoters. Three independent lines of each ZIF-1 transgene from a tissue-specific promoter were scored. The AWC phenotype data of the wild type (row a) and *ajm-1(vy11)* (row b) are the same as those of Figure 1B (rows a and b, respectively). **(B)** Quantification of AWC asymmetry phenotypes in animals expressing *dlg-1::GFP* knock-in and ZIF-1 transgenes, or *GFP::AID::let-413* knock-in and GFP nanobody transgene. Three independent lines of each transgene were scored. **(A, B)** +, heat-shock induction of endogenous protein knockdown at mixed embryonic stages. –, no heat-shock treatment. Animals were scored at the L1 stage. n, total number of animals scored.

To further dissect the timeframe during which *ajm-1* is required for the establishment of AWC asymmetry, we collected and heat-shocked embryos from *ajm-1a-m::ZF1::mNG knock-in*; *hsp-16.2p::zif-1* at 2-hour intervals after egg laying (HAE). The AWC asymmetry phenotype was scored in the L1 or adult stage. Heat shock of the embryos collected at 2-4 HAE, 4-6 HAE, and 6-8 HAE led to a high penetrance (61%-73%) of the 2AWC^OFF^ phenotype (Figure S4B, rows b-d). However, heat shock of the embryos collected at 0-2 HAE and past 8 HAE led to a lower penetrance (22%-38%) of the 2AWC^OFF^ phenotype (Figure S4B, rows a, e, and f). In addition, heat shock of animals at the larval stages did not show an AWC asymmetry defect (Figure S4B, rows g-j). Collectively, these results suggest that *ajm-1* is required for the establishment of AWC asymmetry in embryogenesis, specifically at 2-8 HAE (from the end of gastrulation to about the 3-fold stage) (Figure S4C), but is not required for the maintenance of AWC asymmetry.

### *ajm-1* is required in glial and hypodermal cells to promote AWC^ON^

To further determine the spatial requirement of *ajm-1* in AWC asymmetry, we used *zif-1* transgenes driven by tissue-specific promoters to degrade AJM-1 protein in the tissues of interest. To circumvent embryonic lethality caused by the constitutive degradation of AJM-1 in specific tissues, *zif-1* was expressed in the tissue being examined only at the required embryogenesis stages. To achieve the spatial and temporal control of AJM-1 degradation, the *tissue-specific promoter::loxP::stop::loxP::zif-1* transgene and the integrated *hsp-16.2p::Cre* transgene (a gift from Sander van den Heuvel, Utrecht University) were introduced into the *ajm-1a-m::ZF1::mNG knock-in* strain. Heat-induced Cre expression excises the *loxP::stop::loxP* sequence from the *tissue-specific promoter::loxP::stop::loxP::zif-*1 transgene, thereby expressing ZIF-1 that subsequently degrades AJM-1::ZF1::mNG, in selected tissues at a specific developmental stage.

Mixed-stage embryos of the *ajm-1a-m::ZF1::mNG knock-in*; *hsp-16.2p::Cre*; *tissue-specific promoter::loxP::stop::loxP::zif-*1 were heat-shocked, and AWC phenotypes were examined at the L1 stage. Degradation of AJM-1 by ZIF-1 expressed from the AWC-specific *odr-3p::loxP::stop::loxP::zif-1* or the pan-neuronal *rab-3p::loxP::stop::loxP::zif-1* transgenes led to a low penetrance (7%-19%) of the 2AWC^OFF^ phenotype, compared to 75% in *ajm-1(vy11)* mutants (Figure 6A, data groups b, d, and e). In contrast, degradation of AJM-1 by ZIF-1 expressed from the pan-glial cell-specific *ptr-10p::loxP::stop::loxP::zif-1* or the hypodermal cell-specific *dpy-7p::loxP::stop::loxP::zif-1* transgenes caused a high penetrance (48%-60%) of the 2AWC^OFF^ phenotype (Figure 6A, data groups f and g). Taken together, these results suggest that *ajm-1* is primarily required in non-neuronal tissues, including glial and hypodermal cells, for promoting AWC^ON^. These results are consistent with the non-cell-autonomous role of *ajm-1* in glial and hypodermal cells in AWC asymmetry, as revealed by the rescue experiments (Figure 4).

The *hsp-16.2p::zif-1* extrachromosomal array was also introduced into *ajm-1a,d-m::ZF1::mNG knock-in*, in which *ajm-1b* and *ajm-1c* isoforms were not tagged. This allowed us to further determine whether the isoforms other than *ajm-1b* and *ajm-1c* are required for AWC asymmetry. Heat shock of the *ajm-1a,d-m::ZF1::mNG knock-in; hsp-16.2p::zif-1* embryos resulted in 0%-3% penetrance of the 2AWC^OFF^ phenotype (Figure 6A, data group h), compared to 57-63% penetrance of the 2AWC^OFF^ phenotype from heat shock of the *ajm-1a-m::ZF1::mNG knock-in; hsp-16.2p::zif-1* embryos (Figure 6A, data group c). Consistent with our RNAi data (Figure 1B, rows g-k), these results further support that *ajm-1b* and *ajm-1c* are the major isoforms required to promote AWC^ON^. As *ajm-1b* and *ajm-1c* are the primary isoforms expressed in glial and hypodermal cells (Figure 2F, 2J, and 2L), these results also further support the non-cell-autonomous role for *ajm-1* in promoting AWC^ON^ in these cells (Figures 4 and 5A).

### DLG-1 and LET-413 are required in glial and hypodermal cells to promote AWC^ON^

Previous studies have shown that DLG-1 and LET-413, homologues of the *Drosophila* tumor suppressors Discs large (Dlg) and Scribble, respectively, are required for proper localization of AJM-1 to the apical junctional domain of *C. elegans* epithelia. These junctional components act together to establish and maintain the integrity of apical junctions in epithelial cells (Bossinger, Klebes et al. 2001, Koppen, Simske et al. 2001, McMahon, Legouis et al. 2001, Segbert, Johnson et al. 2004, Lockwood, Lynch et al. 2008). To determine whether DLG-1 or LET-413, like AJM-1, also plays a role in AWC asymmetry, we used a system that combines ZIF-1-mediated protein degradation with a GFP nanobody (Wang, Tang et al. 2017) to degrade endogenous DLG-1::GFP knock-in (Vuong-Brender, Suman et al. 2017, Cohen and Sundaram 2020) or GFP::AID::LET-413 knock-in (Riga, Cravo et al. 2021) in hypodermal and glial cells.

The *tissue-specific promoter::loxP::stop::loxP::GFP nanobody::zif-1* transgene and the integrated *hsp-16.2p::Cre* transgene (a gift from Sander van den Heuvel, Utrecht University) were introduced into *dlg-1::GFP knock-in* strain (Vuong-Brender, Suman et al. 2017, Cohen and Sundaram 2020) or *GFP::AID::let-413 knock-in* strain (Riga, Cravo et al. 2021). Like AJM-1 degradation, degradation of DLG-1 (Figure 6B, data groups c and d) or LET-413 (Figure 6B, data groups e and f) in either the glial or hypodermal cells caused 22%-40% penetrance of the 2AWC^OFF^ phenotype. These results suggest that the apical junction components DLG-1 and LET-413, like AJM-1, play a role in AWC asymmetry. These results also suggest that AJM-1, DLG-1, and LET-413 may function at apical junctions of non-neuronal tissues, including glial and hypodermal cells, to promote AWC^ON^.

### *ajm-1* regulates *slo-1* expression in AWC and other tissues

As the *ajm-1(vy11)* mutation suppressed the *slo-1(ky399gf)* 2AWC^ON^ phenotype, we tested the possibility that *ajm-1* may regulate *slo-1* expression by examining the expression of endogenous *slo-1::GFP knock-in* (Oh, Haney et al. 2017) in wild-type and *ajm-1(vy11)* mutants. Endogenous SLO-1::GFP signals are dim and indistinguishable from the autofluorescence in early-stage embryos (Figure S5A). To overcome this issue, we performed immunostaining with an anti-GFP antibody on wild-type and *ajm-1(vy11)* embryos. An integrated transgene *hlh-16p::H1-wCherry* (Murray, Boyle et al. 2012) was introduced into *slo-1::GFP knock-in* to label the AWC nucleus in embryos. The anti-SLO-1::GFP signal was significantly reduced in AWC and the whole embryo in *ajm-1(vy11)* mutants compared to wild type (Figure S5B). Similar results were observed in wild-type and *ajm-1(vy11)* embryos containing the *slo-1p::2xnls::GFP* transgene, expressing nucleus-localized GFP from a *slo-1* promoter (Alqadah, Hsieh et al. 2016) (Figure S5C). These results suggest that *ajm-1* is required for *slo-1* expression in AWC neurons and other tissues in embryos.

To determine the tissue required for *slo-1* function in specifying the AWC^ON^ subtype, we performed tissue-specific degradation of endogenous SLO-1::GFP knock-in. Loss-of-function allele of either *slo-1* or *slo-2* did not display an AWC asymmetry defect, whereas *slo-1(eg142lf); slo-2(ok2214lf)* double mutants exhibited 100% penetrance of 2AWC^OFF^ phenotype (Figure S6A, data group b) (Alqadah, Hsieh et al. 2016). Therefore, the *GFP nanobody::zif-1* transgene driven by different promoters was introduced into *slo-1::GFP knock-in; slo-2(ok2214lf). slo-1::GFP knock-in* by itself did not show any defects in AWC asymmetry (Figure S6A, data group c), while *slo-1::GFP knock-in; slo-2(ok2214lf)* double mutants exhibited 29% penetrance of 2AWC^OFF^ (Figure S6A, data group d). Heat shock of *slo-1::GFP knock-in; slo-2(ok2214lf); hsp-16.2p::GFP nanobody::zif-1* embryos led to 84%-94% penetrance of 2AWC^OFF^ (Figure S6A, data group e). A similar result was observed in *slo-1::GFP knock-in; slo-2(ok2214lf)* containing the AWC-specific *odr-3p::GFP nanobody::zif-1* transgene, which caused 96%-100% penetrance of 2AWC^OFF^ (Figure S6A, data group f). Degradation of SLO-1 in *slo-1::GFP knock-in; slo-2(ok2214lf),* containing the hypodermal cell or glial cell-specific *GFP nanobody::zif-1* transgene, caused 21-34% penetrance of 2AWC^OFF^, similar to *slo-1::GFP knock-in; slo-2(ok2214lf)* without the transgene (Figure S6A, data groups d and g-i). These results suggest that *slo-1* functions cell autonomously in AWC neurons to promote the AWC^ON^ subtype, consistent with previous findings (Alqadah, Hsieh et al. 2016).

To determine the developmental stage in which *slo-1* functions in AWC asymmetry, embryos of the *slo-1::GFP knock-in; slo-2(ok2214lf); hsp-16.2p::GFP nanobody::zif-1* transgenic lines were collected at different hours after egg laying (HAE). Heat shock of embryos at 2-4 HAE and 4-6 HAE led to 80%-91% penetrance of the 2AWC^OFF^ phenotype (Figure S6B, rows b and c). On the other hand, heat shock of the embryos collected at 0-2 HAE and 6-8 HAE caused 36%-38% penetrance of the 2AWC^OFF^ phenotype (Figure S6B, rows a and d). These results suggest that *slo-1* is required to promote the AWC^ON^ subtype during embryogenesis, specifically from 2-6 HAE, at a stage overlapping with that of *ajm-1* for promoting AWC^ON^ (Figure S4B).

### *ajm-1* antagonizes a mechanosensitive calcium signaling to promote AWC^ON^

Our previous study demonstrated that UNC-2 and EGL-19 voltage-activated calcium channels act redundantly to antagonize the function of SLO-1 and SLO-2 BK channels, thereby promoting the AWC^OFF^ identity (Alqadah, Hsieh et al. 2016). A recent study demonstrated that DEL-1 mechanosensitive DEG/ENaC channel activates EGL-19 to promote dendrite morphogenesis in *C. elegans* (Tao, Coakley et al. 2022). To determine whether *del-1* plays a role in the UNC-2-and EGL-19-mediated calcium signaling pathway to specify AWC^OFF^, we analyzed AWC asymmetry phenotypes in *del-1* mutants and the genetic relationship of *del-1* with other genes in the AWC asymmetry pathway. Both *del-1(ok150)* loss-of-function (lf) (Yemini, Jucikas et al. 2013) and *del-1(wy1014)* gain-of-function (gf) (Tao, Coakley et al. 2022) mutants displayed wild-type AWC asymmetry phenotype (Figure 7, rows a-c). The wild-type AWC asymmetry phenotype was also observed in *egl-19(n582)* reduction-of-function (rf) and *egl-19(ad695gf)* mutants (Figure 7, rows d and e), consistent with previous findings (Bauer Huang, Saheki et al. 2007, Alqadah, Hsieh et al. 2016). As previous studies have shown, *unc-2(lj1lf)* mutants displayed mixed AWC asymmetry phenotypes (Figure 7, row f) (Bauer Huang, Saheki et al. 2007). Similar to *egl-19(n582rf); unc-2(lj1lf)* double mutants (Figure 7, row g) (Bauer Huang, Saheki et al. 2007), *del-1(ok150lf) unc-2(lj1lf)* double mutants displayed a significant enhancement of the *unc-2(lj1lf)* 2AWC^ON^ phenotype (Figure 7, row h). The genetic relationship between *del-1* and *unc-2* is consistent with that between *egl-19* and *unc-2*, suggesting that DEL-1 mechanosensitive DEG/ENaC channel may act in the same pathway as EGL-19 voltage-activated calcium channel to specify AWC^OFF^.

**Figure 7.**
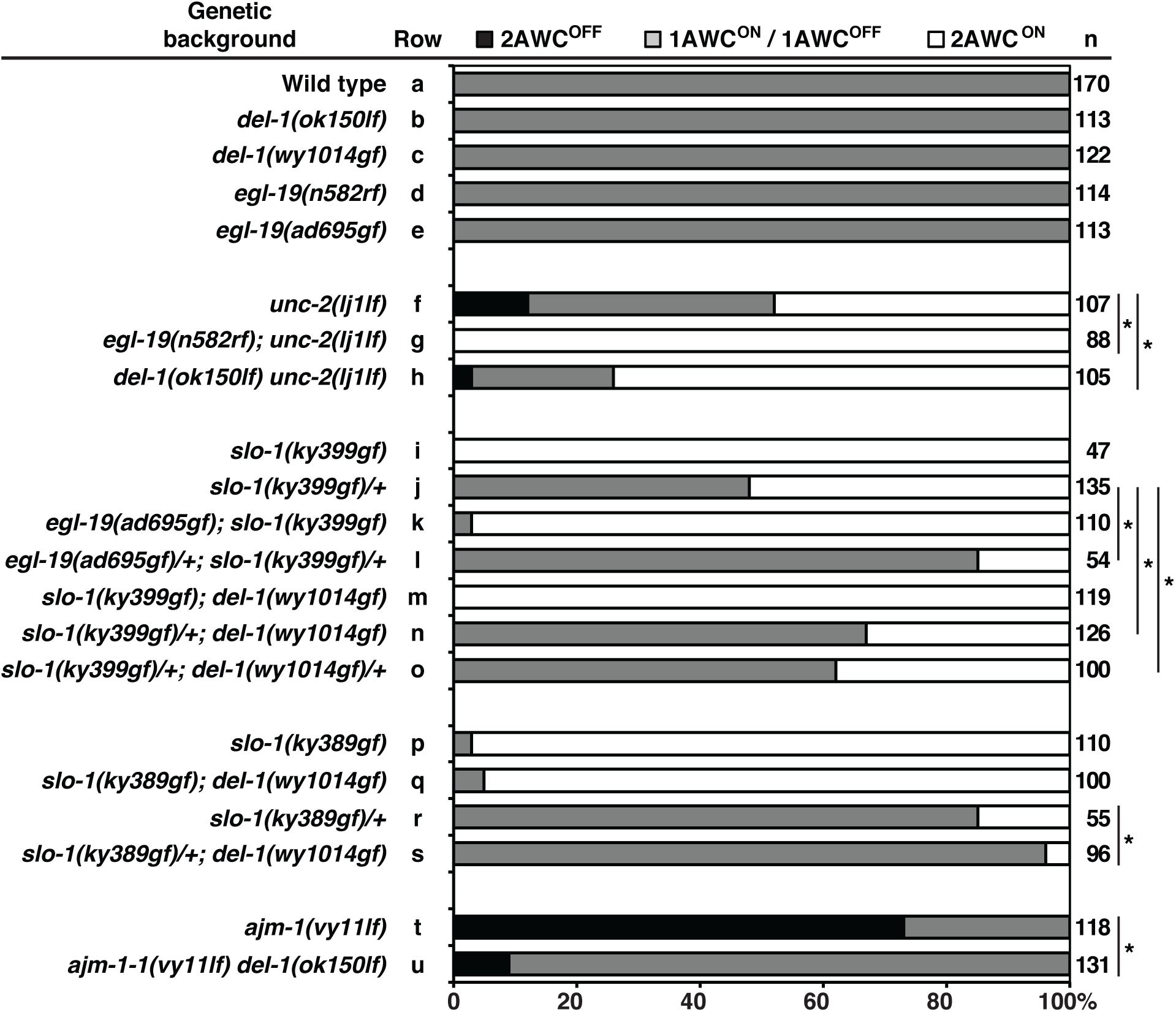
*ajm-1* antagonizes a mechanosensitive calcium signaling to promote AWC^ON^. Quantification of AWC asymmetry phenotypes in single and double mutants of *del-1* (DEG/ENaC channel), *egl-19* (pore-forming α1 subunit voltage-activated L-type calcium channel), *unc-2* (pore-forming α1 subunit voltage-activated P/Q/N-type calcium channel), *slo-1* (voltage- and calcium-activated large conductance BK potassium channel), and *ajm-1* (apical junction molecule). Animals were scored at the adult stage. An asterisk indicates a significant difference between two groups. A *Z-*test was used for statistical analysis. n, total number of animals scored. The AWC phenotype data of the wild type (row a) are the same as those of Figure 1B (row a).

The antagonistic relationship between the EGL-19 voltage-activated calcium channel and the SLO-1 BK potassium channel in AWC asymmetry was previously analyzed using their loss-of-function mutants (Alqadah, Hsieh et al. 2016). To further demonstrate this relationship, double mutants of *egl-19(gf)* and *slo-1(gf)* (Troemel, Sagasti et al. 1999, Davies, Pierce-Shimomura et al. 2003) were analyzed. While *egl-19(ad695gf); slo-1(ky399gf)* double mutants showed a slight reduction in the *slo-1(ky399gf)* 2AWC^ON^ phenotype from 100% to 97% (Figure 7, rows i and k), *slo-1(ky399gf)/+*; *egl-19(ad695gf)*/+ mutants showed significant suppression of the *slo-1(ky399gf)*/+ 2AWC^ON^ phenotype from 52% to 15% (Figure 7, rows j and l). These results further support the antagonistic relationship between EGL-19 and SLO-1 in AWC asymmetry.

To determine whether DEL-1, like EGL-19, antagonizes the function of SLO-1, double mutants of *del-1(gf)* and *slo-1(gf)* were analyzed. Whereas *slo-1(ky399gf); del-1(wy1014gf)* did not show any reduction of the *slo-1(ky399gf)* 2AWC^ON^ phenotype (Figure 7, row m), *slo-1(ky399gf)/+; del-1(wy1014gf)* displayed significant suppression of the *slo-1(ky399gf)* 2AWC^ON^ phenotype from 52% to 33% (Figure 7, row n). In addition, *slo-1(ky399gf)/+; del-1(wy1014gf)/+* showed significant suppression of the *slo-1(ky399gf)* 2AWC^ON^ phenotype from 52% to 38% (Figure 7, row o). Similar experiments were performed using *del-1(wy1014gf)* and *slo-1(ky389gf),* which showed a slightly lower penetrance of the 2AWC^ON^ phenotype than *slo-1(ky399gf)* (Figure 7, rows i and p). While *slo-1(ky389gf); del-1(wy1014gf)* showed only a slight reduction of the *slo-1(ky389gf)* 2AWC^ON^ phenotype from 97% to 95% (Figure 7, rows p and q), *slo-1(ky389gf)/+; del-1(wy1014gf)* showed significant suppression of the *slo-1(ky389gf)*/+ 2AWC^ON^ phenotype from 15% to 4% (Figure 7, rows r and s). Together, these results suggest that, as EGL-19 does, DEL-1 antagonizes SLO-1 function to specify AWC^OFF^. These results also support the above-mentioned notion that DEL-1 mechanosensitive DEG/ENaC channel and EGL-19 calcium channel may act in the same pathway to specify AWC^OFF^.

We also determined the genetic relationship between *ajm-1* and *del-1*. *ajm-1(vy11lf) del-1(ok150lf)* double mutants showed a significant reduction of the *ajm-1(vy11)* 2AWC^OFF^ phenotype from 73% to 9% (Figure 7, rows t and u). This result suggests an antagonistic relationship between *ajm-1* and *del-1* in AWC asymmetry.

Taken together, these genetic data suggest a model in which the AJM-1 apical junction molecule and the SLO-1 BK potassium channel antagonize mechanosensitive calcium signaling, mediated by the DEL-1 mechanosensitive DEG/ENaC channel, the UNC-2 voltage-activated calcium channel, and the EGL-19 voltage-activated calcium channel, to promote AWC^ON^.

To determine the tissues required for *del-1* function in promoting AWC^OFF^, we performed tissue-specific degradation of endogenous DEL-1::GFP knock-in at mixed embryonic stages (Tao, Coakley et al. 2022). While *del-1(lf)* mutants displayed wild-type AWC asymmetry phenotypes (Figures 7, row b, and S7, data group c), they significantly suppressed the *ajm-1(vy11)* 2AWC^OFF^ phenotype (Figures 7, rows t and u, and S7, data groups b and d). Therefore, the *GFP nanobody::zif-1* transgene driven by different promoters was introduced into *ajm-1(vy11) del-1::GFP knock-in* (Figure S7). Degradation of DEL-1::GFP knock-in by heat shock of *ajm-1(vy11) del-1::GFP knock-in; hsp-16.2p::GFP nanobody::zif-1* significantly suppressed the *ajm-1(vy11)* 2AWC^OFF^ phenotype from 47% to 30%-32% (Figure S7, data groups f and g). *ajm-1(vy11) del-1::GFP knock-in* containing the AWC-specific *odr-3p::GFP nanobody::zif-1*, pan-neuronal *rab-3p::GFP nanobody::zif-1*, pan-glial cell-specific *ptr-10p::GFP nanobody::zif-1*, or hypodermal cell-specific *dpy-7p::GFP nanobody::zif-1* transgene also showed significant suppression of the *ajm-1(vy11)* 2AWC^OFF^ phenotype from 57% to 7%-28% (Figure S7, data groups e and h-k). Together, these results suggest that *del-1* may act cell autonomously in AWC neurons and non-cell autonomously in non-AWC cells, including glial and hypodermal cells, to promote AWC^OFF^.

## Discussion

Here, we identify a novel role for AJM-1, an apical junction molecule, in the asymmetric differentiation of one olfactory neuron pair in *C. elegans,* as revealed by an unbiased forward genetic screen. Our findings demonstrate, in addition to its cell-autonomous role, the non-cell-autonomous role of AJM-1 in non-neuronal cells, including glial and hypodermal cells, in antagonizing mechanosensitive calcium signaling mediated by DEL-1 (mechanosensitive DEG/ENaC channel) and voltage-activated calcium channels, UNC-2 (CaV2) and EGL-19 (CaV1), to promote the AWC^ON^ subtype. To the best of our knowledge, this is the first study to implicate the non-cell-autonomous role of apical junction molecules in stochastic cell type choices. Our study also provides insight into the role of mechanical forces in stochastic lateralization of the nervous system.

Our results, together with previous findings, suggest a mechanistic model for the function of *ajm-1* in the stochastic determination of AWC neuron subtypes (Figure 8). During early nerve system development in *C. elegans*, ciliated neurons (including AWC), sheath glial cells, and socket glial cells of the anterior sensilla anchor their processes to the tip of the embryo. As the AWC and glial soma migrate posteriorly during embryogenesis, the anchored dendrites and glial processes stretch out via retrograde extension (Figure 8A) (Heiman and Shaham 2009, Kelley, Yochem et al. 2015, Singhvi and Shaham 2019). This process occurs approximately from the bean stage (about 360 minutes after fertilization) through the 1.5-fold stage (about 460 minutes after fertilization) and possibly until the neurons reach their destination (Figure 8A) (Heiman and Shaham 2009). The timeframe of retrograde extension is consistent with that of *ajm-1* function in AWC asymmetry. In addition, electron microscopy analysis revealed three distinct electron-dense tight junctions between amphid neurons and sheath glia, between sheath and socket glia, and between socket glia and hypodermal cells at the anterior tip of the animal (Ward, Thomson et al. 1975, Perkins, Hedgecock et al. 1986). AJM-1 is localized at these three tight junctions, as shown in this study and previous findings (Figure 8A and 8B) (Nguyen, Liou et al. 2014, Nechipurenko, Olivier-Mason et al. 2016, Low, Williams et al. 2019). The inward-pulling mechanical force exerted by AWC and glial somas during retrograde extension (Kelley, Yochem et al. 2015) may modulate mechanical tension at the three distinct AJM-1-containing tight junctions, thereby regulating the establishment of AWC asymmetry (Figure 8A and 8B).

**Figure 8.**
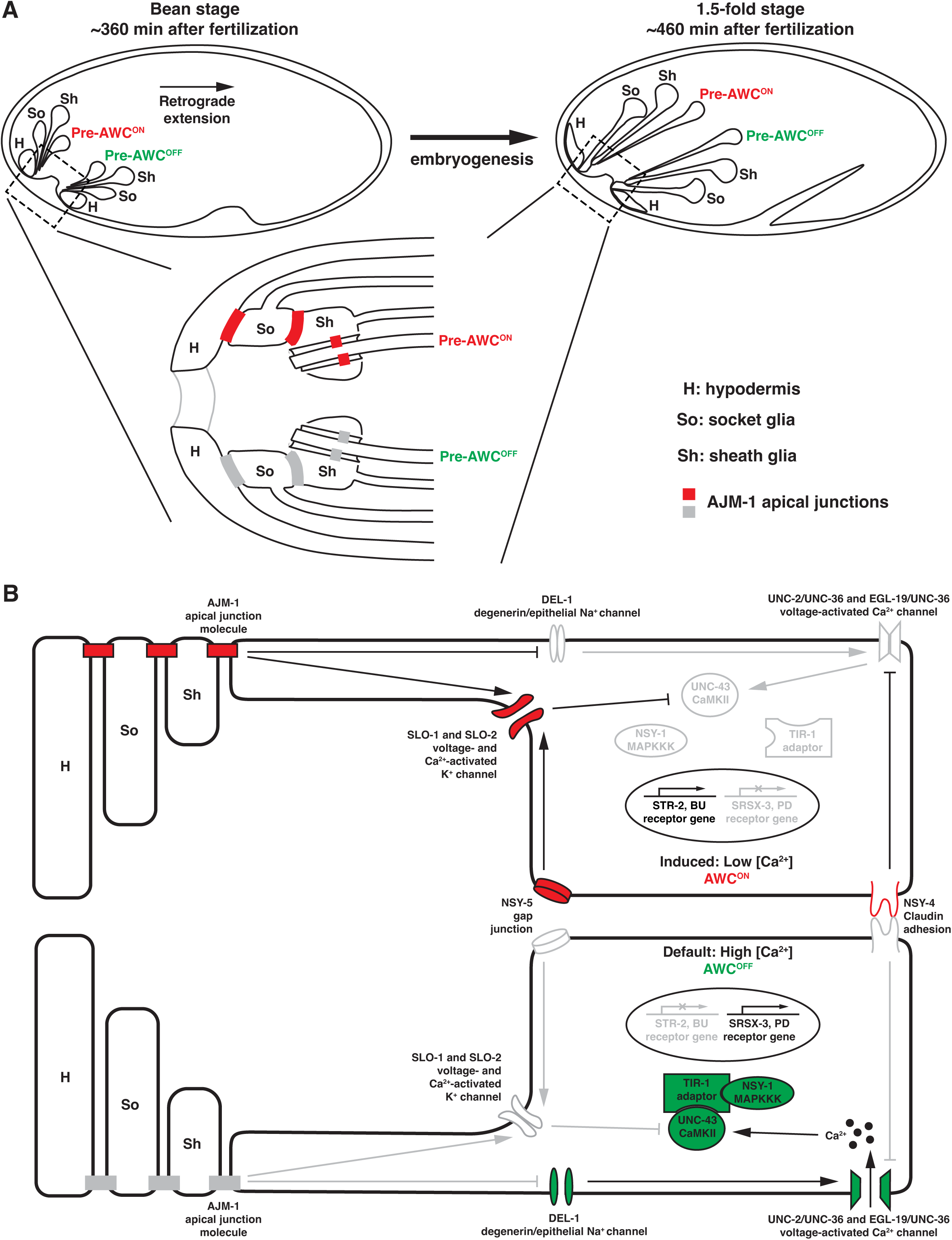
Model of *ajm-1* function in AWC asymmetry. **(A)** Developing AWC neurons, sheath glia, and socket glia migrate posteriorly from anchored dendrites and glial processes via retrograde extension during early embryogenesis. This process may generate mechanical forces sensed by migrating cells. This diagram does not reflect the actual anatomical position of the cells. **(B)** Schematic diagram of *ajm-1* function in promoting AWC^ON^. In the AWC^OFF^ neuron (bottom), DEL-1 mechanosensitive channels activate voltage-activated calcium channels (UNC-2/UNC-36 and EGL-19/UNC-36), leading to the activation of a kinase cascade consisting of UNC-43 (CaMKII), TIR-1 (Sarm1) adaptor protein, and NSY-1 (MAPKKK). This results in the expression of *srsx-3* (AWC^OFF^ marker) and suppression of *str-2* (AWC^ON^ marker). In the AWC^ON^ neuron (Segbert, Johnson et al.), SLO BK potassium channels act to antagonize calcium signaling downstream of the NSY-5 gap junction protein and NSY-4 claudin-like protein. AJM-1 inhibits calcium signaling by promoting SLO-1 expression. Additionally, AJM-1 antagonizes the function of DEL-1 mechanosensitive channels, probably by modulating mechanical forces between non-neuronal and neuronal cells, thereby suppressing voltage-activated calcium channels. Together, these result in the desuppression of *str-2* and the suppression of *srsx-3*. The asymmetric activation of AJM-1 may depend on differences in the mechanical force experienced by the AWC neurons. Red, genes promoting AWC^ON^. Green, genes promoting AWC^OFF^. Grey, genes that are less active or inactive. **(A-B)** Anterior is left, and ventral is down. H, hypodermis; So, socket glia; Sh, sheath glia.

In *C. elegans*, the epidermis is also known as the hypodermis (WormAtlas). In addition to tight junctions, amphid neurons and glial cells share other key features with epithelial cells, including apical-basal polarity. These findings suggest that amphid neurons and glia can be considered continuous with the hypodermis at the anterior tip of the animal (Low, Williams et al. 2019). Epithelial cells are connected by the apical junctional complex, which consists primarily of tight and adherens junctions (Labouesse 2006). In addition, the apical junctional complex regulates mechanical tension by absorbing and dissipating force, which is critical for maintaining the structural integrity of body tissues, including those in the nervous system (Moorthy, Chen et al. 2000, Javier-Torrent, Zimmer-Bensch et al. 2021, Nguyen, Otani et al. 2024). Our previous findings showed that SLO-1 (BK potassium channels) and EGL-19 (L-type voltage-activated calcium channels) have antagonistic and parallel roles in AWC asymmetry, with SLO-1 inhibiting a calcium-activated kinase cascade to promote the AWC^ON^ subtype (Figure 8B) (Alqadah, Hsieh et al. 2016). Our results from this study suggest that AJM-1 acts non-cell autonomously to promote SLO-1 expression and antagonize the function of DEL-1 (mechanosensitive DEG/ENaC channels) in the activation of EGL-19 (Figure 8B). Differential mechanical tension may occur at the three distinct tight junctions: between the amphid neurons and sheath glia, between sheath glia and socket glia, and between socket glia and hypodermal cells, within the two amphid sensilla. This tension may be regulated by the apical junction complex, which includes AJM-1, and is sensed by DEL-1. Consequently, this regulation may lead to differential EGL-19 activity and downstream calcium signaling, ultimately resulting in the asymmetric differentiation of AWC subtypes (Figure 8B).

Mechanical forces have been shown to play a role in brain development and the establishment of left-right body asymmetries. For example, the transient receptor potential (TRP) cationic channel TRPV2 controls axonal outgrowth in developing sensory and motor neurons in response to membrane stretching in mice (Shibasaki, Murayama et al. 2010), while DEG/ENaC channels stimulate dendritic branch growth of sensory neurons in response to mechanical force generated by ligand-receptor interactions in *C. elegans* (Tao, Coakley et al. 2022). In addition, mechanically sensitive heat shock factor 1 is asymmetrically activated by shear stress in the Kupffer’s vesicle, a left-right organizer, during left-right axis establishment in zebrafish embryos (Du, Li et al. 2023). However, the developmental role of mechanical forces in left-right brain development remains unclear. Our study reveals the role of mechanical forces in the stochastic lateralization of the nervous system. Our findings also provide insight into a novel, non-cell-autonomous role for an apical junction molecule in regulating mechanical signaling underlying stochastic brain lateralization.

### Limitations of the Study

We identified a role for the DEL-1 mechanosensitive DEG/ENaC channel in the calcium signaling pathway, mediated by UNC-2 and EGL-19 (voltage-activated calcium channels), to specify AWC^OFF^. Our conclusions are based on the genetic interaction between *del-1* and *unc-2,* which is consistent with that between *egl-19* and *unc-2*, as well as the genetic relationship between *del-1* and *slo-1,* which is consistent with that between *egl-19* and *slo-1*. Since *del-1(gf)* mutants display wild-type AWC asymmetry, we cannot determine whether DEL-1 activates EGL-19 by analyzing *del-1(gf); egl-19(lf)* double mutants. In addition, we cannot directly measure mechanical force at the three tight junctions between amphid neurons and sheath glia, sheath and socket glia, and socket glia and hypodermal cells *in vivo* during embryogenesis due to limitations in available tools.

## Materials and Methods

### Strains and transgenes

The wild-type strain is N2, the *C. elegans* Bristol variety. Strains were maintained using standard methods (Brenner 1974).

### Behavioral assays

Chemotaxis assays were performed, and chemotaxis index was calculated as previously described (Bargmann, Hartwieg et al. 1993, Troemel, Kimmel et al. 1997). 2-butanone (Sigma 360473) and 2,3-pentanedione (Sigma 241962) were diluted 1:1,000 and 1:10,000, respectively, in ethanol. At least 50 animals per genotype were scored for each odor in an independent chemotaxis assay. Assays were repeated three times independently.

### Germline transformation

Germline transformation was performed by injection as described previously (Mello and Fire 1995). DNA was injected into the syncytial gonad of adult hermaphrodites as P_0_. F_1_ animals expressing the co-injected fluorescent marker were selected and singled, and the F_2_ progenies were screened for transgenic lines.

### Generation of *ajm-1::mNG knock-in, ajm-1::TagRFP-T knock-in, ajm-1::GFP11 knock-in*, and *ajm-1::ZF1::mNG knock-in using* Cas9-triggered homologous recombination

The *ajm-1::mNG knock-in, ajm-1::TagRFP-T knock-in,* and *ajm-1::ZF1::mNG knock-in* strains were generated using Cas9-triggered homologous recombination as previously described (Dickinson, Ward et al. 2013, Dickinson, Pani et al. 2015, Dickinson and Goldstein 2016). N2 or *ajm-1(vy11)* animals, cultured at 20 °C, were injected with the construct expressing sgRNA and Cas9, a homology repair template containing mNG/TagRFP-T (or ZF1::mNG) and a SEC cassette, and three co-injection markers [*rab-3p::mCherry* (10 ng/μl), *myo-2p::mCherry* (2.5 ng/μl), and *myo-3p::mCherry* (5 ng/μl)]. The SEC cassette of the homology repair template consists of a hygromycin resistance gene, Cre driven by a heat-shock promoter, and a dominant roller phenotype marker *sqt-1(e1350)*. Injected worms were put on new culture plates (3 worms per plate) and allowed to lay eggs at 25 °C. Three days later, the plates were applied with hygromycin (Gold Biotechnology) at a final concentration of 250 μg/ml and then returned to 25 °C. Three to four days later, animals with a roller phenotype lacking red fluorescent co-injection markers were selected and cloned onto single plates to establish mNG/TagRFP-T^SEC knock-in or ZF1::mNG^SEC knock-in lines. To excise the SEC cassette from the insertion lines, L1 animals were picked for heat shock at 34°C for four hours to induce expression of Cre, then allowed to grow at 20 °C for 5-7 days. Freely moving F_2_ progeny of heat-shocked worms were cloned to establish mNG/TagRFP-T knock-in or ZF1::mNG knock-in lines without the SEC cassette. The correct insertion of mNG/TagRFP-T or ZF1::mNG and the correct excision of SEC were verified by PCR and sequencing.

The *ajm-1::GFP11* knock-in was generated using CRISPR/Cas9-triggered homologous recombination using single-stranded oligodeoxynucleotide donors (ssODN) (Farboud, Severson et al. 2019, Goudeau, Sharp et al. 2021). *muIs253[eft-3p::sfGFP1-10::unc-54 3’UTR Cbr-unc-119(+)] II* animals, cultured at 20 °C, were injected with the construct expressing sgRNA and Cas9, a 194-nucleotide single-stranded oligodeoxyribonucleotide containing GFP11 and *ajm-1* homologous region. Lines were selected and established according to the published protocol (Goudeau, Sharp et al. 2021). muIs253 were removed by genetic crossing to obtain a stand-alone *ajm-1::GFP11* knock-in.

### Tissue-specific visualization of target genes using Split-GFP

The first 10 beta-strands of GFP1-10 (Cabantous, Terwilliger et al. 2005) under the control of tissue-specific promoters were injected into N2 animals. Transgenic lines were selected and genetically crossed with *ajm-1::GFP11* knock-in animals.

### Immunostaining

Antibody staining on mixed-stage *C. elegans* embryos was performed using the “Freeze-Cracking” method (Duerr 2006). For anti-AJM-1 staining, NH27 at 1:100 (DSHB, AB_531819) was used as the primary antibody, and FITC at 1:50 (goat anti-mouse, Jackson ImmunoResearch Lab, 115-095-003) was used as the secondary antibody. For anti-GFP staining, anti-full-length GFP at 1:200 (Clontech, 632593) was used as the primary antibody, and FITC at 1:50 (goat anti-rabbit, Jackson ImmunoResearch Lab, 111-095-003) was used as the secondary antibody.

### Imaging of transgenic animals expressing fluorescent proteins

Animals were anesthetized with 5 mM sodium azide (Sigma) or 7.5 mM levamisole (Sigma) on 2% agarose pads. Images were acquired using a Zeiss Axio Imager M2 microscope, which is equipped with a motorized focus drive, a Zeiss objective EC Plan-Neofluar 63x/1.40 Oil DIC M27, a Zeiss objective EC Plan-Neofluar 40x/1.30 Oil DIC M27, a Piston GFP bandpass filter set (41025, Chroma Technology), a TRITC filter set (41002c, Chroma Technology), a Zeiss Apotome system, a Hamamatsu digital camera C11440, and the Zeiss imaging software ZEN (2012 blue edition SP2).

### Quantification of fluorescence intensity

Transgenic animals expressing fluorescent markers were imaged using the same exposure time in each set of experiments. In Figure S5B, z-stack images of 1.5-fold stage embryos showing GFP expression were projected into a single image, and then the fluorescence intensity of the whole embryo was measured using the ZEISS Zen software. In Figure S5C, a single focal plane with the brightest GFP fluorescence in the AWC nucleus was selected from a stack of images for the measurement of fluorescence intensity using the ZEISS Zen software.

### Genetic mosaic analysis

Genetic mosaic analysis was performed and analyzed as described previously (Chuang, Vanhoven et al. 2007). Transgenic animals containing unstable extrachromosomal arrays were passed for at least 6 generations before analysis. The co-injection marker *odr-1p::DsRed*, expressed in both AWC and AWB neurons, was used to determine the presence or absence of the extrachromosomal transgene in AWC and AWB cell lineages.

### Targeted degradation of endogenous proteins

Degradation of proteins tagged with ZF1 zinc-finger domain is induced upon expression of the E3 ubiquitin ligase substrate-recognition subunit ZIF-1 (Armenti, Lohmer et al. 2014). Degradation of proteins tagged with GFP is induced upon expression of GFP nanobody fused with ZIF-1 (Wang, Tang et al. 2017). *zif-1* or *GFP nanobody::zif-1* controlled by tissue-specific promoter or heat-shock promoter *hsp-16.2p* was injected into *ZF1::mNG* or *GFP knock-in* animals. To temporally control the tissue-specific degradation of target proteins, a *let-858* terminator region flanked by *loxP* sites was constructed in between tissue-specific promoters and *zif-1* or *GFP nanobody::zif-1*. These constructs were injected into GFP knock-in strains, which also contain an integrated *hsp-16.2p::Cre*. Larvae or embryos from transgenic lines were heat-shocked at 34 ℃ for 30 min and then transferred to 20 ℃. Animals were allowed to develop to the adult stage, and the percentage of animals with the 2AWC^OFF^ phenotype was analyzed.

## Supplemental Information Appendix

Supplemental information includes supplemental materials and methods, supplemental references, supplemental figure legends, and supplemental figures S1-S7.

## Data availability

All data discussed in the paper will be available to readers.

## Acknowledgments

We thank Alex Boyanov, Oliver Hobert, Dan Dickinson, Bob Goldstein, Daniel Shaye, Hongkyun Kim, Jeffery Simske, Maxwell G. Heiman, Shai Shaham, Mike Boxem, Shen Kang, Sander van den Heuvel, and WormBase for assistance, strains, reagents, protocols, and/or databases. We also thank Peter Sahyouni for comments on the manuscript. Some strains were provided by the Caenorhabditis Genetics Center (CGC), which is funded by NIH Office of Research Infrastructure Programs (P40 OD010440). This work was supported by a Whitehall Foundation Research Award (to C.-F.C.), an Alfred P. Sloan Research Fellowship (to C.-F.C.), and the National Institutes of Health Grant (5R01GM098026- 05 to C.-F.C.).

## Author Contributions

Conceptualization, C.-F.C.; Methodology, R.X. and C.-F.C.; Investigation, R.X., J.Y., SY., X.W., and C.-F.C.; Formal Analysis, R.X., J.Y., SY., and C.-F.C.; Writing – Original Draft, R.X. and C.-F.C.; Writing – Review and Editing, R.X., E.L., and C.-F.C; Funding Acquisition, C.-F.C.; Supervision, C.-F.C.

## Supplemental Information

### Supplemental Materials and Methods

#### Strains and transgenes

##### Mutants

*eri-1(mg366)* IV; *lin-15b(n744)* X (Sieburth, Ch’ng et al. 2005)

*slo-1(ky399gf)* V (Troemel, Sagasti et al. 1999)

*slo-1(ky389gf)* V (Troemel, Sagasti et al. 1999)

*slo-1(eg142lf)* V (Kim, Pierce-Shimomura et al. 2009)

*slo-2(ok2214lf)* V (*C. elegans* knockout consortium)

*ajm-1(vy11lf)* X (this study)

*ajm-1(ok160lf)* X (Koppen, Simske et al. 2001)

*unc-2(lj1lf)* X (Tam, Mathews et al. 2000)

*egl-19(n582rf)* IV (Lee, Lobel et al. 1997)

*egl-19(ad695gf)* IV (Lee, Lobel et al. 1997)

*del-1(ok150lf)* X (*C. elegans* knockout consortium)

*del-1(wy1014gf)* X (Tao, Coakley et al. 2022)

##### Fluorescent protein knock-in strains

*ajm-1a-m::ZF1::mNG knock-in* (Figures 2, 6, S2, S3, S4)

*ajm-1(vy359[ajm-1a-m::ZF1::mNG^3xFLAG])* X (this study)

Parent strain of *ajm-1(vy359)*: *ajm-1(vy356[ajm-1a-m::ZF1::mNG^SEC^3xFLAG])* X (this study)

*ajm-1a,c-m::mNG knock-in* (Figures 2, S2)

*ajm-1(vy189[ajm-1a,c-m::mNG^3xFLAG])* X (this study)

Parent strain of *ajm-1(vy189)*: *ajm-1(vy175[ajm-1a,c-m::mNG^SEC^3xFLAG])* X (this study)

*ajm-1a,d-m::ZF1::mNG knock-in* (Figures 2, 6, S2, S4)

*ajm-1(vy319[ajm-1a,d-m::ZF1::mNG^3xFLAG])* X (this study)

Parent strain of *ajm-1(vy319)*: *ajm-1(vy318[ajm-1a,d-m::ZF1::mNG^SEC^3xFLAG])* X (this study)

*mNG::ajm-1b,c,d,f knock-in* (Figures 2, S2)

*ajm-1(vy314[mNG^3xFLAG::ajm-1b,c,d,f])* X (this study)

Parent strain of *ajm-1(vy314)*: *ajm-1(vy313[mNG^SEC^3xFLAG::ajm-1b,c,d,f])* X (this study)

*ajm-1(vy11)a,c-m::mNG knock-in* (Figure 2)

*ajm-1(vy340[ajm-1(vy11)a,c-m::mNG^3xFLAG])* X (this study)

Parent strain of *ajm-1(vy340)*: *ajm-1(vy338[ajm-1(vy11)a,c-m::mNG^SEC^3xFLAG])* X (this study)

*ajm-1a-m::GFP11 knock-in* (Figures 2, S2)

*ajm-1(vy434[ajm-1a-m::GFP11])* X (this study)

*ajm-1a,c-m::TagRFP-T knock-in* (Figures 3, S2)

*ajm-1(vy438[ajm-1a,c-m::TagRFP-T^3xFLAG])* X (this study)

Parent strain of *ajm-1(vy438)*: *ajm-1(vy435[ajm-1a,c-m::TagRFP-T^SEC^3xFLAG])* X (this study)

*ajm-1(vy11)a,c-m::TagRFP-T knock-in* (Figure 3)

*ajm-1(vy440[ajm-1(vy11)a,c-m::TagRFP-T^3xFLAG])* X (this study)

Parent strain of *ajm-1(vy440)*: *ajm-1(vy439[ajm-1(vy11)a,c-m::TagRFP^SEC^3xFLAG])* X (this study)

*GFP::unc-70 knock-in* (Figure 3)

*unc-70(cas962[GFP::unc-70])* V (Jia, Li et al. 2019)

*dlg-1::GFP knock-in* (Figures 6 and S4)

ML2615 *dlg-1(mc103[dlg-1::GFP])* X (Cohen, Sparacio et al. 2020)

*GFP::loxP::AID::let-413 knock-in* (Figures 6 and S4)

BOX466 *let-413(mib81[GFP::loxP::AID::let-413])* X (Riga, Cravo et al. 2021)

*slo-1::GFP knock-in* (Figures S5 and S6)

*slo-1(cim105[slo-1::GFP])* V (Oh, Haney et al. 2017)

*del-1::GFP knock-in* (Figure S7)

*del-1(wy1149[del-1::GFP]* X (Tao, Coakley et al. 2022)

##### Integrated transgenes

| Integrated transgenes | Figures |
| --- | --- |
| *vyIs68 [str-2p::TagRFP; srsx-3p::GFP]* III (Cochella, Tursun et al. 2014) | 1 |
| *kyIs140 [str-2p::GFP; lin-15(+)]* I (Troemel, Sagasti et al. 1999) | 1, 4, 5, 6, 7,<br>S1, S2, S4,<br>S6, S7 |
| *kyIs323 [str-2::GFP; ofm-1::GFP]* II | 1 |
| *jcIs1* [*ajm-1::GFP* + *unc-29(+)* + *rol-6(su1006)*] IV (Koppen, Simske et al. | 2 |
| *vyIs56* [*odr-1p::TagRFP*] III (Cochella, Tursun et al. 2014) | 2, S1 |
| *oxTi973* [*eft-3p::GFP::2xNLS::tbb-2 3' UTR* + *HygroR*] V (Frokjaer-Jensen, Davis et al. 2014) | 3 |
| *vyIs96* [*str-2p::TagRFP*] V (this study) | 6, S4, S7 |
| *heSi157* [*hsp-16.2p::Cre*] X (Gift from Sander van den Heuvel, Utrecht University) | 6, S4, S6 |
| *otIs264* [*ceh-36p::tagRFP*] (Patel, Tursun et al. 2012) | S1 |
| *stIs10544* [*hlh-16::H1-wCherry::let-858 3' UTR*] (Murray, Boyle et al. 2012) | S5 |
| *vyIs97* [*str-2p::TagRFP*] IV (this study) | S6 |

##### Extrachromosomal arrays

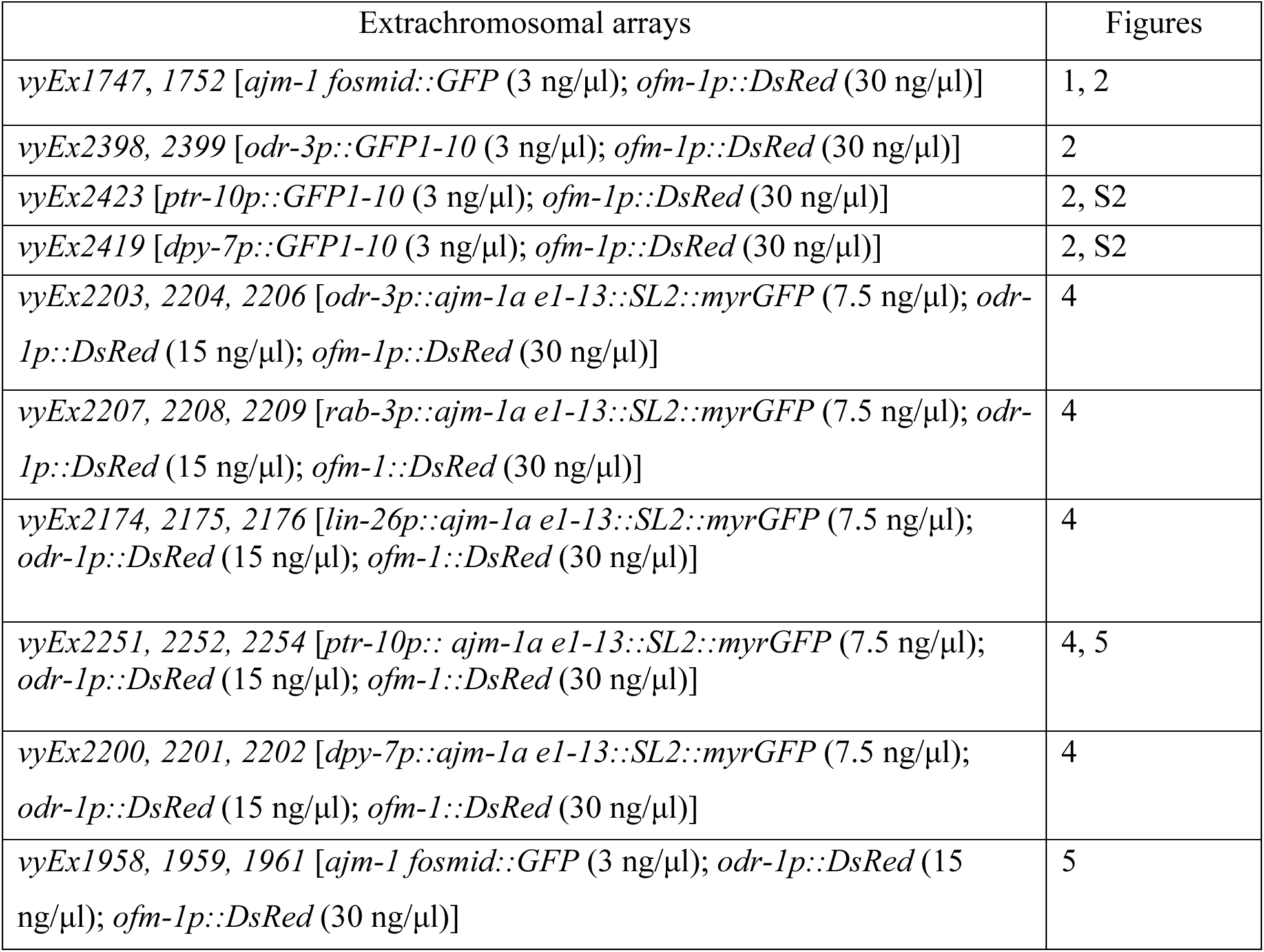
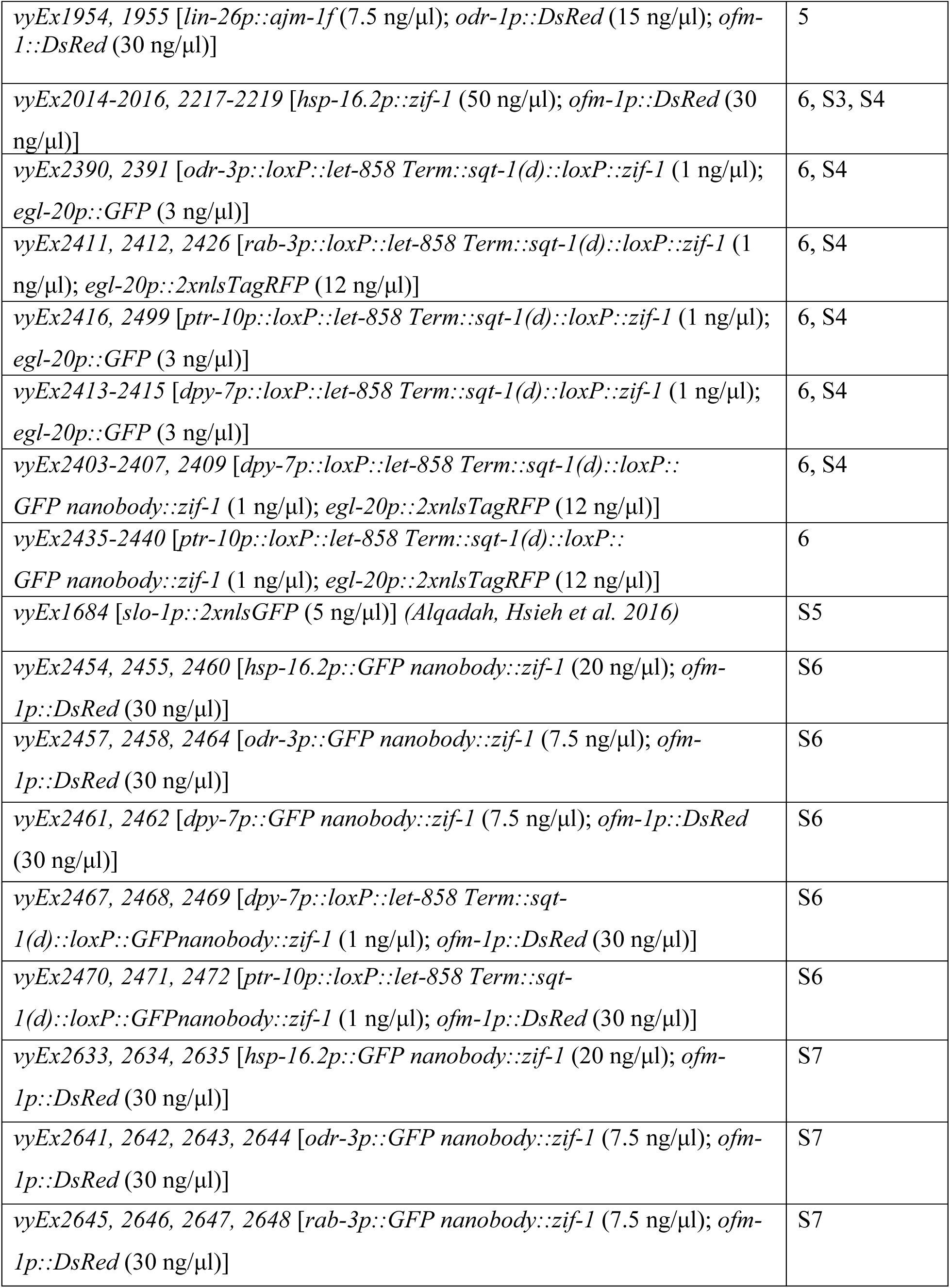
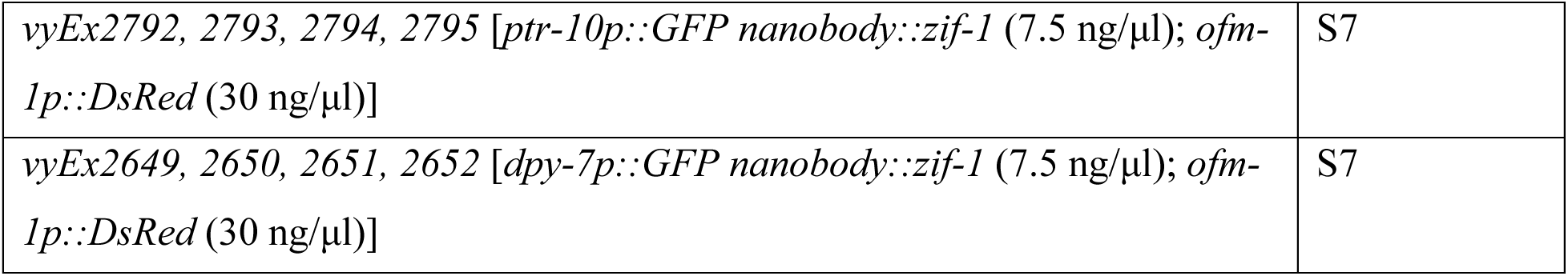

#### Plasmid construction

*ajm-1b 662bp RNAi* was generated by subcloning a 662 bp fragment (F: GTCTGGCATGAAAACAGAAATAATTTCTG; B: TGAAAATGTGTACGTGTAGGTTGTTC) containing the last 11 bp *ajm-1b* exon 9 and 651 bp of *ajm-1b 3’ UTR* into the pAD12 vector (Dillin, Crawford et al. 2002).

*ajm-1c RNAi* was generated by subcloning a 397 bp fragment (F: GTTGTTGCATTATAAGCCACTTTTCAATC; B: GAATTTGATAGTTTTTATTTATTCAAAATTTGT TCATACATAAAC) containing the last 15 bp of the *ajm-1c* exon 13 and 382 bp of the *ajm-1c 3’ UTR* into the pAD12 vector (Dillin, Crawford et al. 2002).

*dlg-1 RNAi* was generated by subcloning a 3462 bp fragment (F: AAGCGCATAAAGCGATAGAAAATGTG; B: CTAATGACGTGGCACCCAAATTG) spanning *dlg-1* exon 2 to 12 into the pAD12 vector (Dillin, Crawford et al. 2002).

*let-413* RNAi was generated by subcloning a 4204 bp fragment (F: CGATCAACTTCATATATCGCCACAATG; B: TTAGGAAGCATACACGGGAGTAC) covering all *let-413* exons into the pAD12 vector (Dillin, Crawford et al. 2002).

*eft-3p::Cas9::U6p::ajm-1a-m sgRNA* was generated by inserting *ajm-1a-m sgRNA* (CATCCTCAACCCATACGTTA) into the *pDD162 eft-3p::Cas9::empty sgRNA* vector (Dickinson, Ward et al. 2013, Dickinson, Pani et al. 2015, Dickinson and Goldstein 2016) (Addgene #47549 from Bob Goldstein’s lab) using the Q5 site-directed mutagenesis kit (New England Biolabs).

*eft-3p::Cas9::U6p::ajm-1a,c-m sgRNA* was generated by inserting *ajm-1a,c-m sgRNA* (ACAGGAAACGTCGAACCGAC) into the *pDD162 eft-3p::Cas9::empty sgRNA* vector (Dickinson, Ward et al. 2013, Dickinson, Pani et al. 2015, Dickinson and Goldstein 2016) (Addgene #47549 from Bob Goldstein’s lab) using the Q5 site-directed mutagenesis kit (New England Biolabs).

*eft-3p::Cas9::U6p::ajm-1a,d-m sgRNA* was generated by inserting *ajm-1a,d-m sgRNA* (GATTCTGAATGGCTTCGTAG) into the *pDD162 eft-3p::Cas9::empty sgRNA* vector (Dickinson, Ward et al. 2013, Dickinson, Pani et al. 2015, Dickinson and Goldstein 2016) (Addgene #47549 from Bob Goldstein’s lab) using the Q5 site-directed mutagenesis kit (New England Biolabs).

*eft-3p::Cas9::U6p::ajm-1b,c,d,f sgRNA* was generated by inserting *ajm-1b,c,d,f sgRNA* (GTCACAGAAAAGTCGTCCTC) into the *pDD162 eft-3p::Cas9::empty sgRNA* vector (Dickinson, Ward et al. 2013, Dickinson, Pani et al. 2015, Dickinson and Goldstein 2016) (Addgene #47549 from Bob Goldstein’s lab) using the Q5 site-directed mutagenesis kit (New England Biolabs).

*ajm-1a-m::ZF1^mNG ^SEC^3xFLAG* was constructed by PCR amplifying a 1500 bp *ajm-1* 5’ homology region (F: AGAAGAGAGAAAAACAATGGAACGTCTTGAAAG; B: GGATCCGTTTCTATTCAAATCATTCGGGTTG) and a 1531 bp *ajm-1* 3’ homology region (F: AGCAGGTGAGACAACTTTCAGCATG; B: CCATACACTAAACATTTCAGCGTGTCTTACTG), and subcloning into the pDS676 ZF1^mNG^SEC^3xFLAG vector (a gift from Daniel Shaye) containing a 111 bp of ZF1 sequence encoding a PIE-1 C-C-C-H type zinc-finger domain tag (Armenti, Lohmer et al. 2014). The nucleotide of the PAM site of the construct was mutagenized to generate a silent mutation from CATCCTCAACCCATACGTTA <u>CGG</u> to CATCCTCAACCCATACGTTt <u>CtG</u> using the Q5 site-directed mutagenesis kit (New England Biolabs).

*ajm-1a,c-m::mNG ^SEC^3xFLAG* was constructed by PCR amplifying a 572 bp *ajm-1* 5’ homology region (F: TTTCTATTTCCGGTCCTCGCAC; B: TAATGCAACAACCTGTCGGTTC) and a 545 bp *ajm-1* 3’ homology region (F: TAAGCCACTTTTCAATCCCAATTACC; B: GAGGACTTCTAATGCTTTAGGGAGC), and subcloning into the pDD240 mNG^SEC^3xFLAG vector (Dickinson, Ward et al. 2013, Dickinson, Pani et al. 2015, Dickinson and Goldstein 2016). The nucleotide of the PAM site of the construct was mutagenized to generate a silent mutation from ACAGGAAACGTCGAACCGAC <u>AGG</u> to ACAGGAAACGTCGAACCGAC <u>AaG</u> using the Q5 site-directed mutagenesis kit (New England Biolabs).

*ajm-1a,d-m::ZF1^mNG ^SEC^3xFLAG* was constructed by PCR amplifying a 1500 bp *ajm-1* 5’ homology region (F: TGCATACTAGCCGCACTTC; B: GACATTGTGTGCAAATTCTGAGAAG) and a 1518 bp *ajm-1* 3’ homology region (F: TAGATATAAATTCATCTTCATTTTTATTATGAGCC; B: GAGTCAATGACCACCTCCG), and subcloning into the pDS676 mNG^SEC^3xFLAG vector (a gift from Daniel Shaye) containing ZF1 zinc-finger domain tag (Armenti, Lohmer et al. 2014). The nucleotide of the PAM site of the construct was mutagenized to generate a silent mutation from GATTCTGAATGGCTTCGTAG <u>CGG</u> to GATTCTGAATGGCTTCGTAG <u>CGt</u> using the Q5 site-directed mutagenesis kit (New England Biolabs).

*ajm-1b,c,d,f::mNG ^SEC^3xFLAG* was constructed by PCR amplifying a 1469 bp *ajm-1* 5’ homology region (F: TTCTCCATCATCATTTTGCC; B: ACAATTCTGAAAAAAAAACCTTTAAAATTTTC) and a 1428 bp *ajm-1* 3’ homology region (F: ATGGACATAGAAAATCTACAATCG; B: TCCTGTCAGTTTTACACATTAAG), and subcloning into the pDD268 mNG^SEC^3xFLAG vector (Dickinson, Ward et al. 2013, Dickinson, Pani et al. 2015, Dickinson and Goldstein 2016). The nucleotide of the PAM site of the construct was mutagenized to generate a silent mutation from GTCACAGAAAAGTCGTCCTC <u>CGG</u> to GTCACAGAAAAGTCGTCCTC <u>CaG</u> using the Q5 site-directed mutagenesis kit (New England Biolabs).

*ajm-1a-m::GFP11 ssODN* was a 194-nucleotide single-stranded oligodeoxyribonucleotide ordered from IDT; the complete sequences are as follows (*ajm-1* homology region in uppercase letters):

AATGACGAGAACAACCGTGATCGTCTTTACAACCCGAATGATTTGAATAGAAACGGATCCGgaggaggatcccgtgaccacatggtccttcatgagtatgtaaatgctgctgggattacaggcggaggttctAGCAGGTGAGACAACTTTCAGCATGGTCTTCAATGCCCAACATAATACAAAAGTTGTTGCAG *ajm-1a,c-m::TagRFP-T ^SEC^3xFLAG* was constructed by PCR amplifying a 1514 bp *ajm-1* 5’ homology region (F: ACTCTGAATTCCGAACCAC; B: TAATGCAACAACCTGTCGGTTCG) and a 1500 bp *ajm-1* 3’ homology region (F: TAAGCCACTTTTCAATCCCAATTACC; B: ACACTTGAACATTGGAATATAAGTTGTTTACC), and subcloning into the pDD284 TagRFP-T^SEC^3xFLAG vector (Dickinson, Ward et al. 2013, Dickinson, Pani et al. 2015, Dickinson and Goldstein 2016). The nucleotide of the PAM site of the construct was mutagenized to generate a silent mutation from ACAGGAAACGTCGAACCGAC <u>AGG</u> to ACAGGAAACGTCGAACCGAC <u>AaG</u> using the Q5 site-directed mutagenesis kit (New England Biolabs).

*odr-3p::GFP1-10* was generated by subcloning a 648 bp GFP region containing the first 10 β-strands into a vector containing 2677 bp of the *odr-3* promoter (Roayaie, Crump et al. 1998).

*ptr-10p::GFP1-10* was generated by subcloning a 648 bp GFP region containing the first 10 β-strands into a vector containing 300 bp of the *ptr-10* promoter (Yoshimura, Murray et al. 2008).

*dpy-7p::GFP1-10* was generated by subcloning a 648 bp GFP region containing the first 10 β-strands into a vector containing 217 bp of the *dpy-7* promoter (Gilleard, Barry et al. 1997).

*odr-3p::ajm-1a e1-13::SL2::myrGFP* was generated by subcloning a 3155 bp of *ajm-1a* cDNA region spanning from the first ATG of *ajm-1a* exon 1 to the first 144 bp of *ajm-1a* exon 13 into a vector containing a 2677 bp of *odr-3* promoter (Roayaie, Crump et al. 1998) and 238 bp of SL2 trans-splicing sequence followed by 891 bp of myrGFP sequence.

*rab-3p::ajm-1a e1-13::SL2::myrGFP* was generated by subcloning a 3155 bp of *ajm-1a* cDNA region spanning from the first ATG of *ajm-1a* exon 1 to the first 144 bp of *ajm-1a* exon 13 into a vector containing a 4383 bp of *rab-3* promoter (Stefanakis, Carrera et al. 2015) and 238 bp of SL2 trans-splicing sequence followed by 891 bp of myrGFP sequence.

*lin-26p::ajm-1a e1-13::SL2::myrGFP* was generated by subcloning a 3155 bp of *ajm-1a* cDNA region spanning from the first ATG of *ajm-1a* exon 1 to the first 144 bp of *ajm-1a* exon 13 into a vector containing a 5000 bp of *lin-26* promoter (Labouesse, Sookhareea et al. 1994, Labouesse, Hartwieg et al. 1996, Landmann, Quintin et al. 2004) and 238 bp of SL2 trans-splicing sequence followed by 891 bp of myrGFP sequence.

*lin-26p::ajm-1f* was generated by subcloning 5715 bp of *ajm-1f* cDNA into a vector containing 5000 bp of the *lin-26* promoter (Labouesse, Sookhareea et al. 1994, Labouesse, Hartwieg et al. 1996, Landmann, Quintin et al. 2004).

*ptr-10p::ajm-1a e1-13::SL2::myrGFP* was generated by subcloning a 3155 bp of *ajm-1a* cDNA region spanning from the first ATG of *ajm-1a* exon 1 to the first 144 bp of *ajm-1a* exon 13 into a vector containing a 300 bp of *ptr-10* promoter (Yoshimura, Murray et al. 2008) and 238 bp of SL2 trans-splicing sequence followed by 891 bp of myrGFP sequence.

*dpy-7p::ajm-1a e1-13::SL2::myrGFP* was generated by subcloning a 3155 bp of *ajm-1a* cDNA region spanning from the first ATG of *ajm-1a* exon 1 to the first 144 bp of *ajm-1a* exon 13 into a vector containing a 217 bp of *dpy-7* promoter (Gilleard, Barry et al. 1997) and 238 bp of SL2 trans-splicing sequence followed by 891 bp of myrGFP sequence.

*hsp-16.2p::zif-1* was generated by cloning 1744 bp of *zif-1* genomic DNA (Armenti, Lohmer et al. 2014) into a vector containing 394 bp of the *hsp-16.2* promoter (Bacaj and Shaham 2007).

*odr-3p::loxP::let-858 Term::sqt-1(d)::loxP::zif-1* was generated by cloning 1744 bp of *zif-1* genomic DNA (Armenti, Lohmer et al. 2014) after a *loxP*-flanking region that contains 381 bp of the let-858 terminator region and a *sqt-1* dominant marker (Dickinson, Pani et al. 2015) into a vector containing 2677 bp of the *odr-3* promoter (Roayaie, Crump et al. 1998).

*rab-3p::loxP::let-858 Term::sqt-1(d)::loxP::zif-1* was generated by cloning 1744 bp of *zif-1* genomic DNA (Armenti, Lohmer et al. 2014) after a *loxP*-flanking region that contains a 381 bp let-858 terminator region and a *sqt-1* dominant marker (Dickinson, Pani et al. 2015) into a vector containing 4383 bp of the *rab-3* promoter (Stefanakis, Carrera et al. 2015).

*ptr-10p::loxP::let-858 Term::sqt-1(d)::loxP::zif-1* was generated by cloning 1744 bp of *zif-1* genomic DNA (Armenti, Lohmer et al. 2014) after a *loxP*-flanking region that contains a 381 bp let-858 terminator region and a *sqt-1* dominant marker (Dickinson, Pani et al. 2015) into a vector containing 300 bp of the *ptr-10* promoter (Yoshimura, Murray et al. 2008).

*dpy-7p::loxP::let-858 Term::sqt-1(d)::loxP::zif-1* was generated by cloning 1744 bp of *zif-1* genomic DNA (Armenti, Lohmer et al. 2014) after a *loxP*-flanking region that contains 381 bp of the let-858 terminator region and a *sqt-1* dominant marker (Dickinson, Pani et al. 2015) into a vector containing 217 bp of the *dpy-7* promoter (Gilleard, Barry et al. 1997).

*hsp-16.2p::GFP nanobody::zif-1* was generated by cloning a 351 bp GFP nanobody fused with 1744 bp of *zif-1* genomic DNA (Wang, Tang et al. 2017) into a vector containing 394 bp of the *hsp-16.2* promoter (Bacaj and Shaham 2007).

*dpy-7p::loxP::let-858 Term::sqt-1(d)::loxP::GFP nanobody::zif-1* was generated by cloning 351 bp GFP nanobody fused with 1744 bp of *zif-1* genomic DNA (Wang, Tang et al. 2017) after a *loxP*-flanking region that contains a 381 bp let-858 terminator region and a *sqt-1* dominant marker (Dickinson, Pani et al. 2015) into a vector containing 217 bp of the *dpy-7* promoter (Gilleard, Barry et al. 1997).

*ptr-10p::loxP::let-858 Term::sqt-1(d)::loxP::GFP nanobody::zif-1* was generated by cloning 351 bp GFP nanobody fused with 1744 bp of *zif-1* genomic DNA (Wang, Tang et al. 2017) after a *loxP*-flanking region that containing a 381 bp let-858 terminator region and a *sqt-1* dominant markers (Dickinson, Pani et al. 2015) into a vector containing 300 bp of the *ptr-10* promoter (Yoshimura, Murray et al. 2008).

*odr-3p::GFP nanobody::zif-1* was generated by cloning a 351 bp GFP nanobody fused with 1744 bp of *zif-1* genomic DNA (Wang, Tang et al. 2017) into a vector containing 2677 bp of the *odr-3* promoter (Roayaie, Crump et al. 1998).

*rab-3p::GFP nanobody::zif-1* was generated by cloning a 351 bp GFP nanobody fused with 1744 bp of *zif-1* genomic DNA (Wang, Tang et al. 2017) into a vector containing 4383 bp of the *rab-3* promoter (Roayaie, Crump et al. 1998).

*ptr-10p::GFP nanobody::zif-1* was generated by cloning a 351 bp GFP nanobody fused with 1744 bp of *zif-1* genomic DNA (Wang, Tang et al. 2017) into a vector containing 300 bp of the *ptr-10* promoter (Roayaie, Crump et al. 1998).

*dpy-7::GFP nanobody::zif-1* was generated by cloning 351 bp GFP nanobody fused with 1744 bp of *zif-1* genomic DNA (Wang, Tang et al. 2017) into a vector containing 217 bp of the *dpy-7* promoter (Roayaie, Crump et al. 1998).

## Supplemental Figure Legends

**Figure S1.**
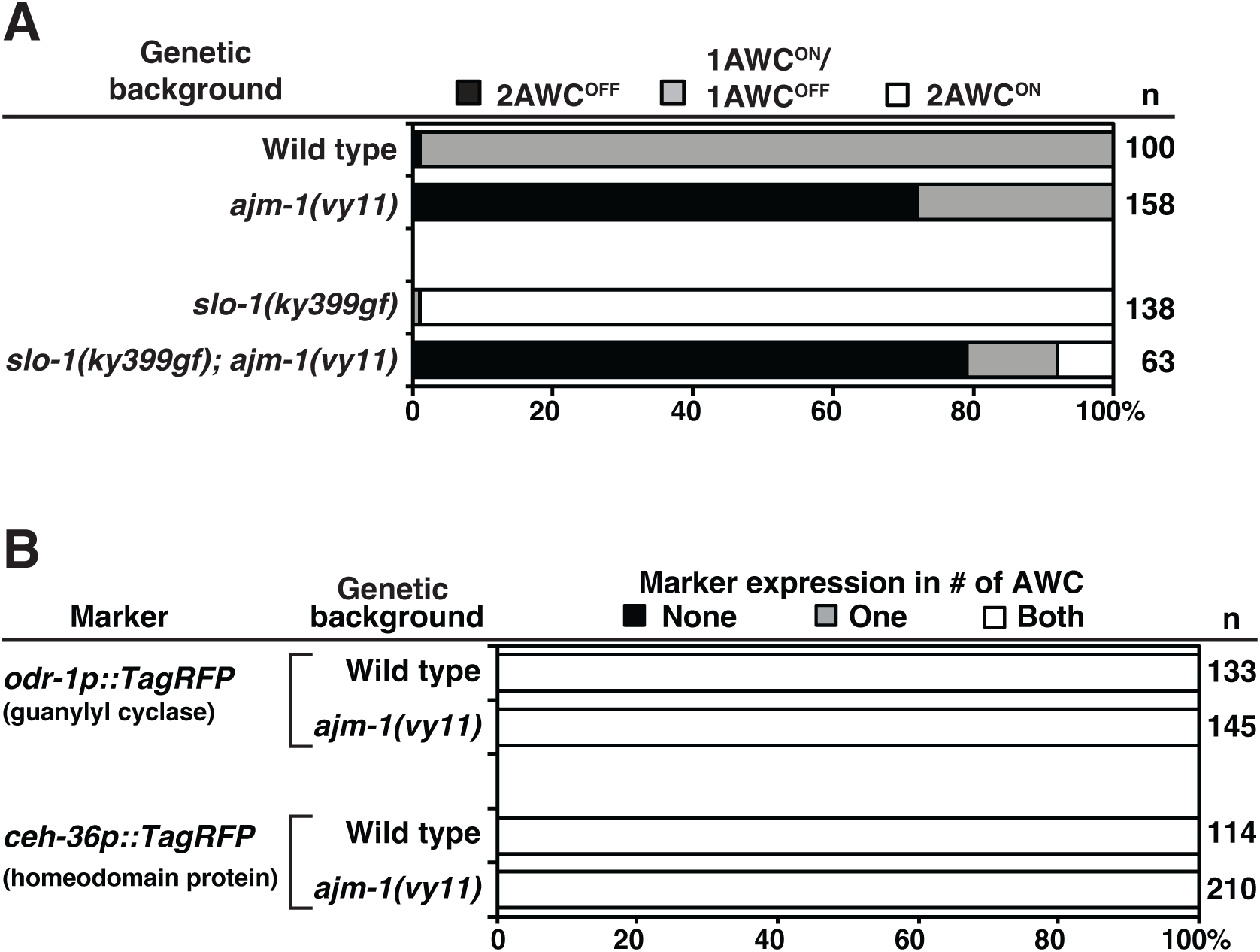
Genetic interactions of *ajm-1(vy11)* with *slo-1(gf)* in AWC asymmetry. **(A)** Double mutant analysis of *ajm-1(vy11)* with *slo-1(ky399gf)* mutants. Animals were scored at the adult stage. n, total number of animals scored. **(B)** Quantification of general AWC identity phenotypes in wild type and *ajm-1(vy11)* mutants. Animals were scored at the adult stage. n, total number of animals scored.

**Figure S2.**
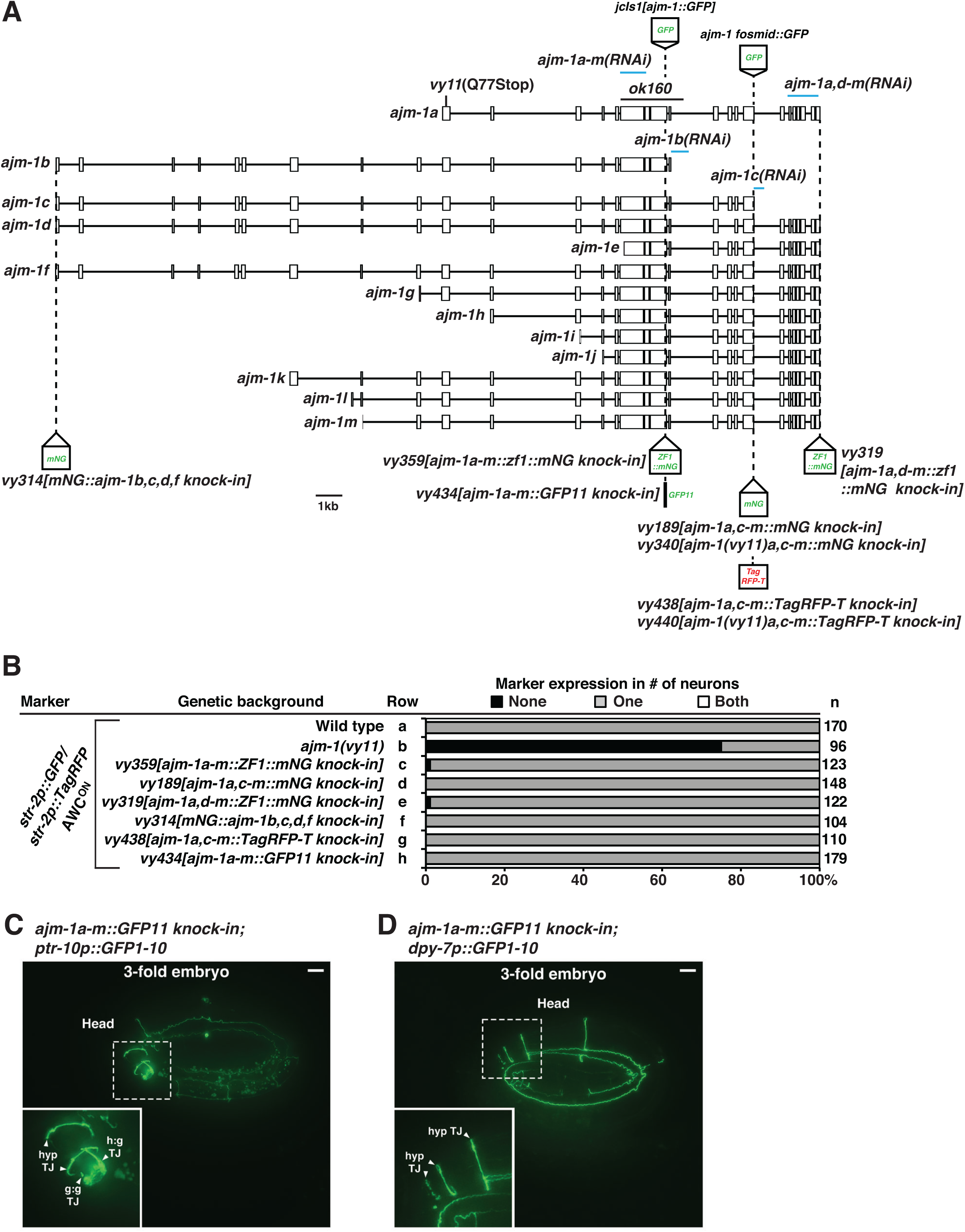
Gene structure and analyses of *ajm-1* knock-in with various fluorescent tags. (**A**) Genomic structure of all *ajm-1* isoforms a-m with the location of CRISPR/Cas9 knock-in tags, mNG, TagRFP-T, or ZF1::mNG. RNAi target regions of different *ajm-1* isoforms are indicated in blue lines. (**B**) Expression of the AWC^ON^ markers in adults. n, total number of animals scored. The AWC phenotype data of the wild type (row a) and *ajm-1(vy11)* (row b) are the same as those of Figure 1B (rows a and b, respectively). (**C**, **D**) Tissue-specific visualization of endogenous AJM-1 protein expression in 3-fold stage embryos using split GFP in specific tissues. GFP11 is knocked in all AJM-1 isoforms a-m. GFP1-10 is under the control of a tissue-specific promoter expressed in pan-glial cells (C) or hypodermal cells (D). Scale bars, 5 μm

**Figure S3.**
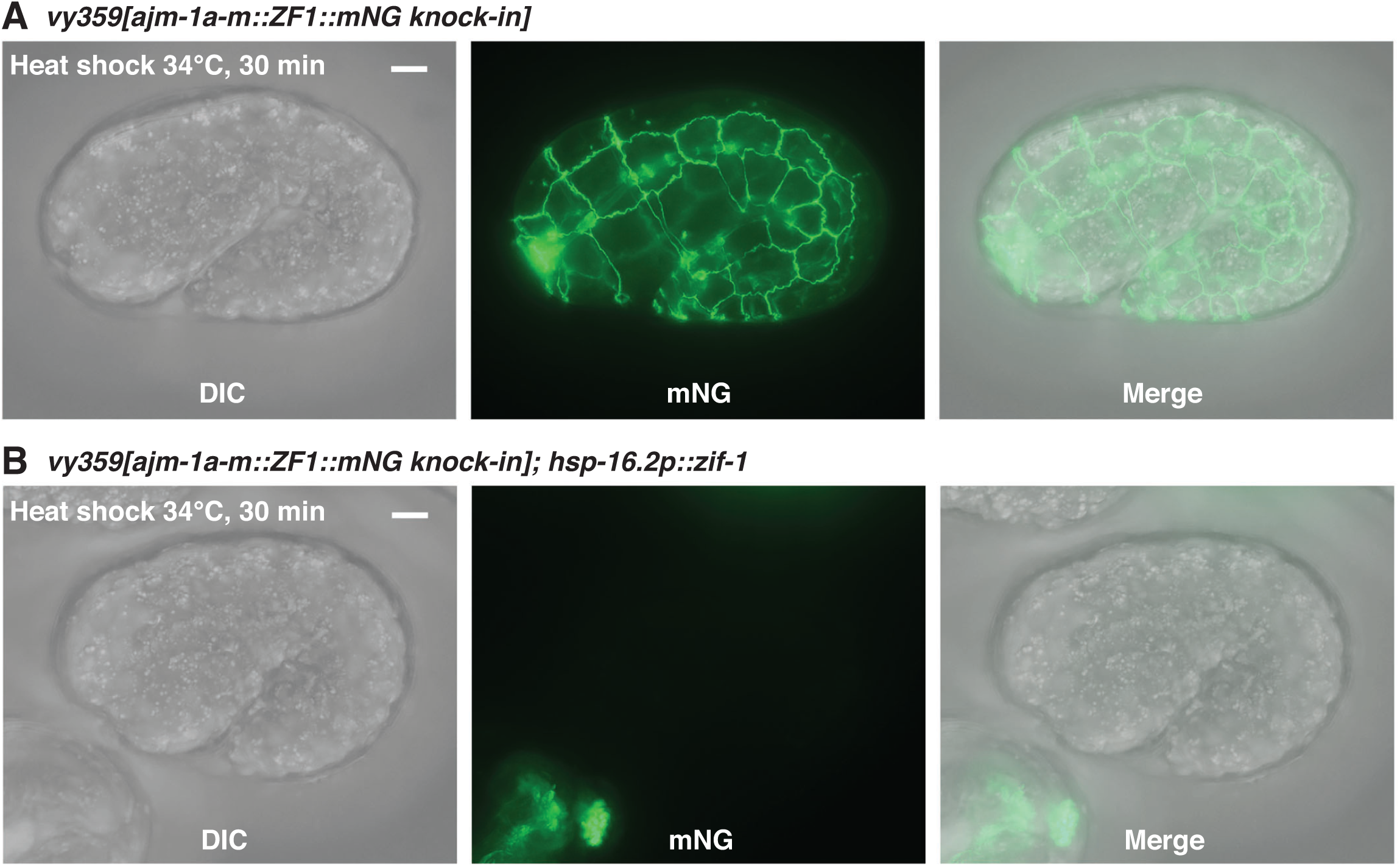
AJM-1a-m::ZF1::mNG is degraded upon heat-shock treatment. (**A**, **B**) Images of *ajm-1(vy359[ajm-1a-m::ZF1::mNG knock-in])* 1.5-fold embryo without (A) and with (B) *hsp-16.2p::zif-1* transgene after heat-shock treatment at 34 ℃ for 30 min. Images were taken 1.5 hrs after the heat-shock treatment finished. Scale bars, 5 μm.

**Figure S4.**
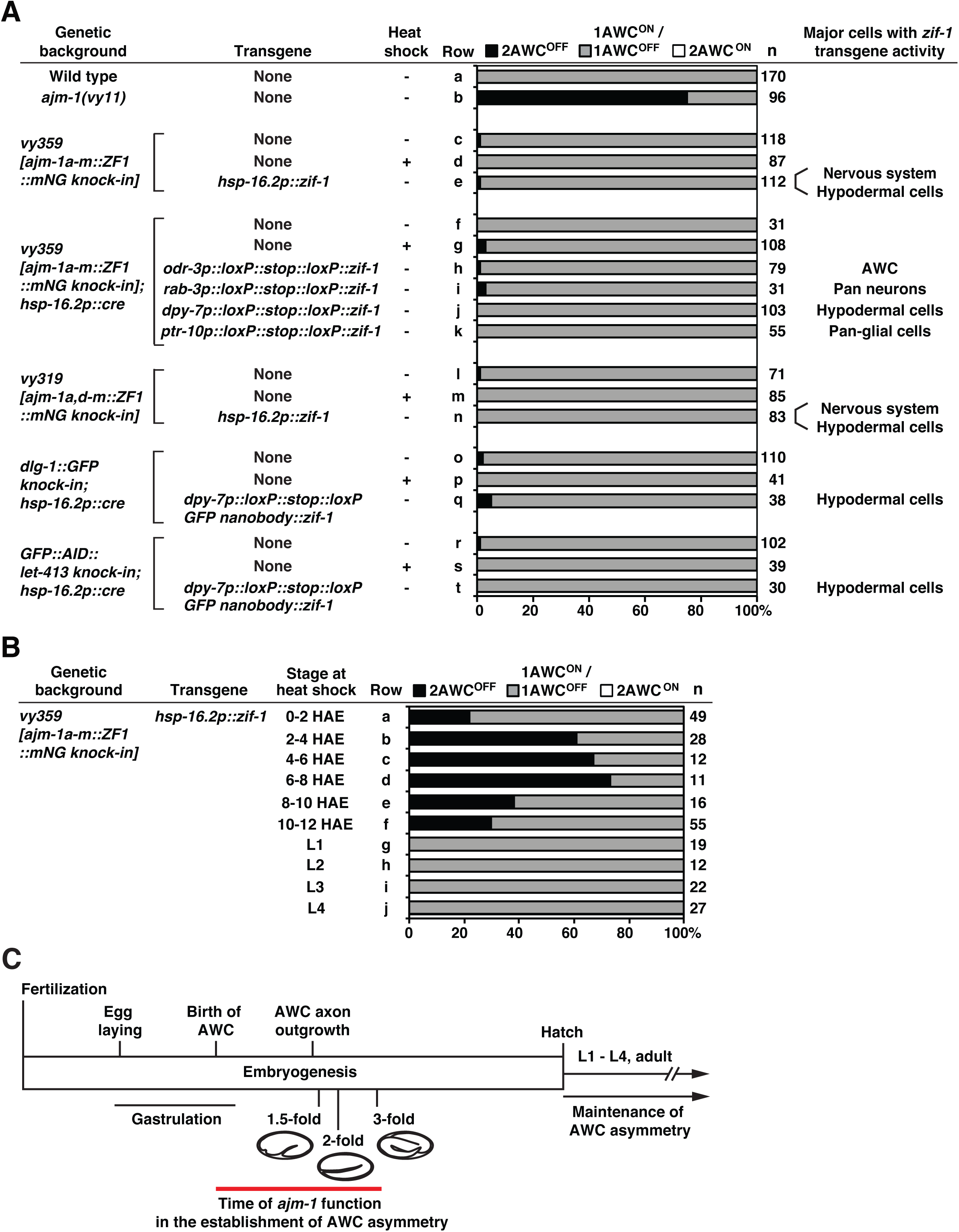
*ajm-1* acts embryonically to promote AWC^ON^. **(A)** Quantification of AWC asymmetry phenotypes in *ajm-1a-m::ZF1::mNG knock-in* (all AJM-1 isoforms tagged), *ajm-1a,d-m::ZF1::mNG knock-in* (AJM-1 isoform b and c not tagged), *dlg-1::GFP knock-in,* and *GFP::AID::let-413 knock-in* animals with heat-shock treatment in the absence of a tissue-specific knockdown transgenic array, or without heat shock in the presence of a transgenic array. +, heat-shock induction of endogenous protein knockdown at mixed embryonic stages. –, no heat-shock treatment. Adult animals from one or two independent lines (combined) of each transgene were scored. n, total number of animals scored. The AWC phenotype data of the wild type (row a) and *ajm-1(vy11)* (row b) are the same as those of Figure 1B (rows a and b, respectively). **(B)** Quantification of AWC asymmetry phenotypes in animals with knockdown of endogenous AJM-1 protein isoforms a-m containing *ZF1::mNG knock-in* at different developmental stages. Transgenic line expressing ZIF-1 (the substrate-binding subunit of the E3 ligase) from the heat-inducible promoter, *hsp-16.2p*, was scored. Embryos and larvae at different stages were collected and heat-shocked. HAE, hours after egg laying. Animals were scored at the adult stage. L1, first-stage larva. L2, second-stage larva. L3, third-stage larva. L4, fourth-stage larva. n, total number of animals scored. (**C**) Timeline of *C. elegans* developmental events at 22°C. Eggs are laid about 150 minutes after fertilization. Gastrulation occurs 140-330 minutes after fertilization. AWC neurons are born at about 300 min, and their axons extend at about 450 min after fertilization. Embryos reach the 1.5-, 2-, and 3-fold stages at about 460 min, 490 min, and 550 min after fertilization, respectively. Animals hatch at about 840 minutes after fertilization. AWC asymmetry is established during embryogenesis and is maintained throughout the animal’s life span.

**Figure S5.**
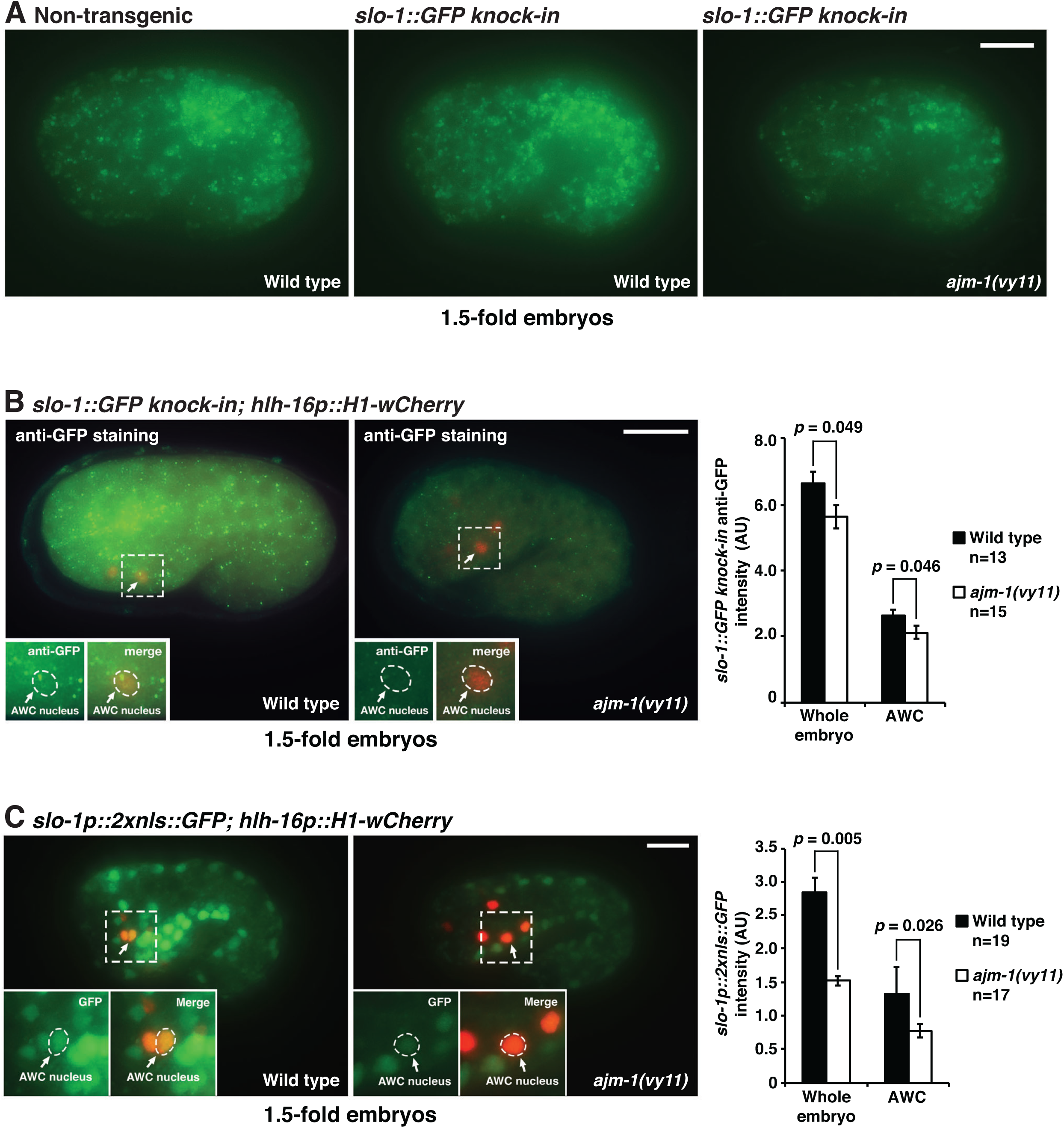
*slo-1* expression is decreased in *ajm-1(vy11)* mutants. **(A)** Images of 1.5-fold stage embryos from wild type (non-transgenic), *slo-1(cim105[slo-1::GFP knock-in])*, and *slo-1(cim105[slo-1::GFP knock-in]); ajm-1(vy11).* Scale bar, 10 μm. **(B)** Left panels: Images of *slo-1(cim105[slo-1::GFP knock-in])* 1.5-fold stage embryos immunostained with an anti-GFP antibody in wild type and *ajm-1(vy11)*. Right panel: Quantification of anti-GFP immunostaining intensity in whole embryos and AWC. A stack of focal planes was projected with maximum intensity and compared for fluorescence intensity. **(C)** Left panels: Images of wild-type and *ajm-1(vy11)* 1.5-fold stage embryos expressing *slo-1p::2xnlsGFP* in the nucleus. Right panel: Quantification of *slo-1p::2xnlsGFP* expression in whole embryos and AWC. The single focal plane with the brightest GFP fluorescence in the AWC at the top of the acquired image stacks was selected and compared for fluorescence intensity of the AWC nucleus. **(B, C)** Insets, indicated by dashed boxes, are magnified by 2-fold. Arrows indicate the AWC nuclei. The AWC nucleus expressing *hlh-16p::H1-wCherry* is outlined with dashed lines. Scale bar, 10 μm. n, total number of animals quantified. Student’s *t*-test was used for statistical analysis. Error bars, standard error of the mean. AU, arbitrary unit.

**Figure S6.**
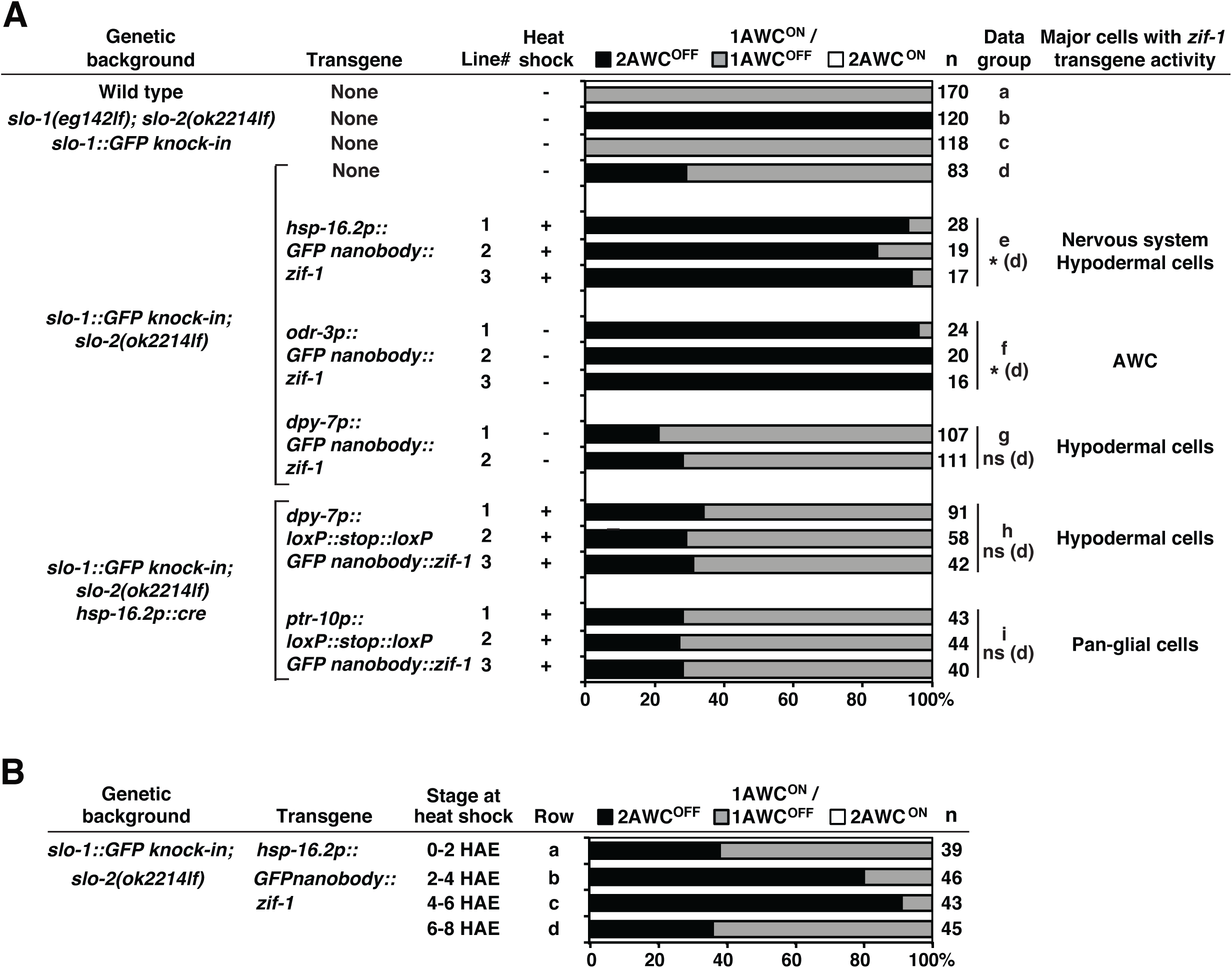
*slo-1* acts cell autonomously to promote the AWC^ON^ subtype. **(A)** Quantification of AWC asymmetry phenotypes in animals with knockdown of endogenous SLO-1 protein containing *GFP* knock-in, in specific tissues. Two or three independent lines expressing GFP nanobody::ZIF-1 from the heat-inducible promoter, *hsp-16.2p*, hypodermis-specific promoter, *dpy-7p*, or AWC-specific promoter, *odr-3p*, were scored. +, heat-shock induction of endogenous protein knockdown at mixed embryonic stages. –, no heat shock treatment. Animals were scored at the adult stage. n, total number of animals scored. Statistical comparisons between individual data groups e-i versus d were determined by a *Z*-test. Asterisks indicate comparisons that are different at *p* < 0.05. ns, not significant. The AWC phenotype data of the wild type (row a) are the same as those of Figure 1B (row a). **(B)** Quantification of AWC asymmetry phenotypes in animals with knockdown of endogenous SLO-1 protein containing *GFP* knock-in at different developmental stages. Transgenic line expressing ZIF-1 from the heat-inducible promoter, *hsp-16.2p*, was scored. Mixed-stage embryos were heat-shocked at various stages. HAE, hours after egg laying. Animals were scored at the adult stage. n, total number of animals scored.

**Figure S7.**
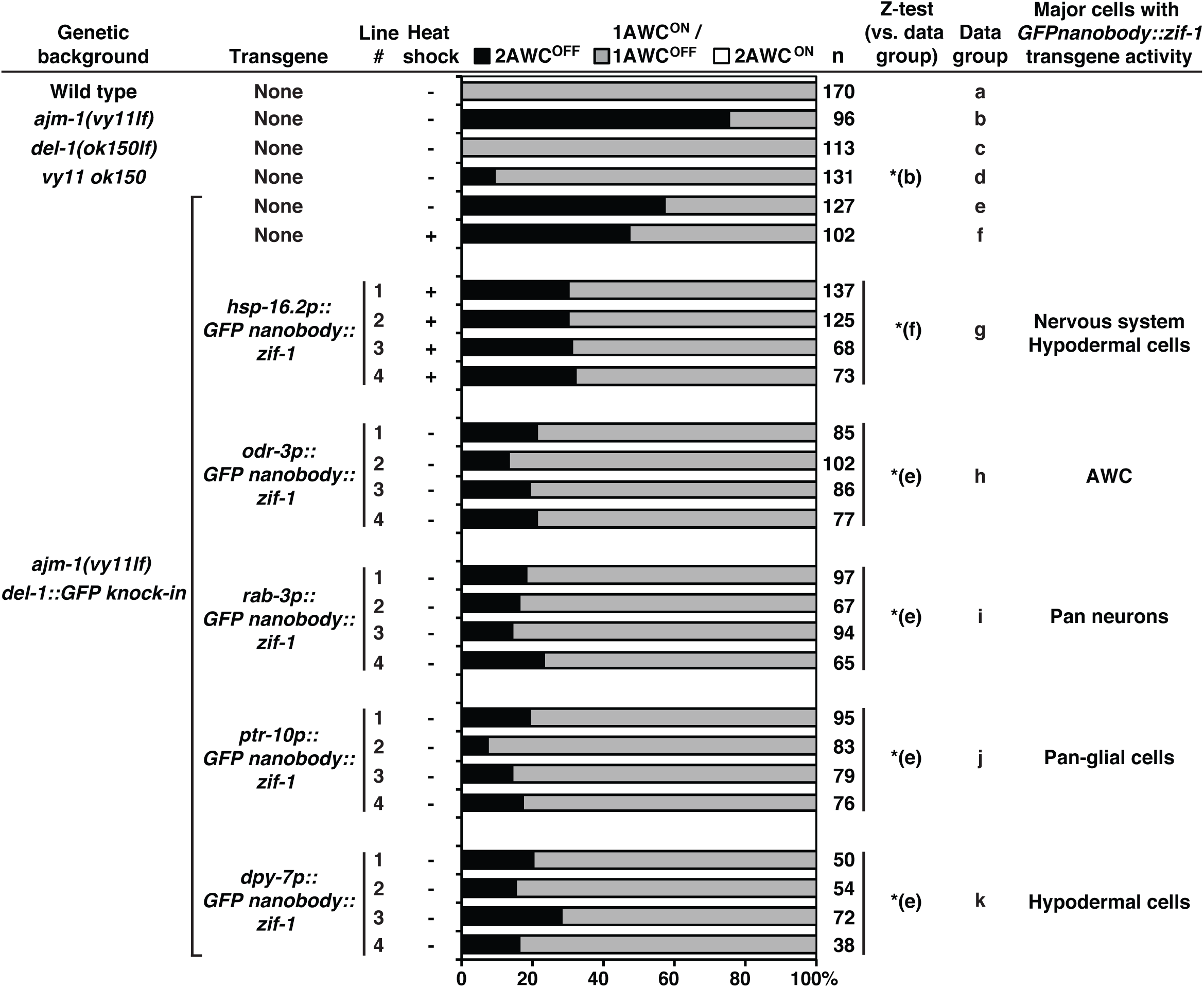
*del-1* acts both cell and non-cell autonomously in AWC asymmetry. Quantification of AWC asymmetry phenotypes in animals with knockdown of endogenous DEL-1 protein containing *GFP* knock-in in specific tissues. Four independent lines expressing GFP nanobody::ZIF-1 from the heat-inducible promoter, *hsp-16.2p*, AWC-specific promoter, *odr-3p*, pan-neuronal promoter, *rab-3p*, pan-glial cell-specific promoter, *ptr-10p*, or hypodermis-specific promoter, *dpy-7p*, were scored. +, heat-shock induction of endogenous protein knockdown at mixed embryonic stages. –, no heat shock treatment. Animals were scored at the adult stage. n, total number of animals scored. Statistical comparisons were made by a *Z*-test. Asterisks indicate comparisons that are different at *p* < 0.05. ns, not significant. The AWC phenotype data of the wild type (row a) and *ajm-1(vy11)* (row b) are the same as those of Figure 1B (rows a and b, respectively). The AWC phenotype data of the *del-1(ok150lf)* (row c) and *vy11 ok150* (row d) are the same as those of Figure 7 (rows b and u, respectively).

